# Estimating the probable cause of recurrence in Plasmodium vivax malaria: relapse, reinfection or recrudescence?

**DOI:** 10.1101/505594

**Authors:** Aimee R Taylor, James A Watson, Cindy S Chu, Kanokpich Puaprasert, Jureeporn Duanguppama, Nicholas P J Day, Francois Nosten, Daniel E Neafsey, Caroline O Buckee, Mallika Imwong, Nicholas J White

## Abstract

**Background:** Relapse of *Plasmodium vivax* infection is the main cause of vivax malaria in many parts of Asia. However at the individual patient level, recurrence of a blood stage infection following treatment within the endemic area can be either a relapse (from dormant liver-stage parasites), a recrudescence (blood-stage treatment failure), or a reinfection (following a new mosquito inoculation). Each requires a different prevention strategy, but previously they could not be distinguished reliably. Time-of-event and genetic data provide complimentary information about the cause of *P. vivax* recurrence, but the optimum approach to genotyping and analysis remains uncertain.

**Methods:** Individual-level data from two large drug trials in acute vivax malaria patients (Vivax History: VHX; Best Primaquine Dose: BPD) conducted on the Thailand-Myanmar border with follow-up of one year were pooled (n=1299). A total of 710 isolates from both acute and recurrent *P. vivax* episodes were genotyped using 3-9 highly polymorphic microsatellite markers. These pooled data were analyzed using a novel population statistical model incorporating an assessment of genetic relatedness, treatment drug administered, and the time-to-recurrence.

**Results:** 99% of genotyped recurrences in individuals who did not receive primaquine (n=365) were estimated to be relapses. In comparison, 14% of genotyped recurrences (n=121) were estimated to be relapses following high-dose supervised primaquine. By comparing episodes across individuals (90194 comparisons), the false-positive rate of relapse identification using genetic data alone was estimated to be 2.2%. We estimated the true failure rate after high-dose primaquine (7mg/kg total dose) to be 2.6% in this epidemiological context, substantially lower the reinfection unadjusted estimate of 12%. Simulation studies show that 9 highly polymorphic microsatellite markers suffice to discriminate between recurrence states. Drug exposures reflected by plasma carboxy-primaquine concentrations were not predictive of treatment failure, but did identify non-adherence.

**Conclusion:** Using this novel statistical model, relapse of *P. vivax* malaria could be distinguished reliably from reinfection. This showed that in this population supervised high-dose primaquine could avert up to 99% of relapses. In low transmission settings, microsatellite genotyping combined with time-to-event data can accurately discriminate between the different causes of recurrent *P. vivax* malaria.

**Author summary:** One hundred years ago, *Plasmodium vivax*, the most globally diverse cause of human malaria, was present across most of the old-world, throughout tropical and temperate climes. Its main evolutionary advantage over *Plasmodium falciparum*, responsible for the most deadly type of human malaria, is its ability to stay dormant in the liver, emerging weeks to years later causing recurring illness and continuing transmission. The dormant liver-stage parasites are called hypnozoites. A recurrent infection can either be hypnozoite-derived (a relapse), a blood-stage treatment failure (recrudescence), or from a new mosquito bite (a reinfection). At a population level, each requires a different preventative strategy but no widely applicable methodology existed to discriminate between the three possible states: relapse, recrudescence and reinfection. Parasite genetic data alone cannot resolve the different possibilities. We developed a novel probabilistic framework which uses both epidemiological and genetic data to determine the most likely cause of vivax recurrence at the individual level. We applied this method to the largest available pooled clinical trial data from Southeast Asia incorporating information from highly polymorphic microsatellite markers, drug treatments received and time-to-recurrent illness. This analysis provides a tool for estimating probabilities of relapse, recrudescence and reinfection and, importantly, shows that hypnozoiticidal high-dose primaquine radical cure is much more effective than previously thought.

## Introduction

### Background

*Plasmodium vivax* is the most geographically widespread *Plasmodium* spp. that causes human malaria, with an estimated 2.5 billion people at risk of infection [1]. According to the 2017 WHO World Malaria Report, there were an estimated 6-11 million cases of vivax malaria in 2016. Many of these illnesses were in the WHO Southeast Asia region and the majority can be ascribed to relapse [2-4].

*P. vivax* and *P. ovale* malarias are characterized by their ability to relapse, which results from the activation of dormant liver-stage parasites called hypnozoites. Multiple relapses can follow a single mosquito inoculation [5]. Two distinct phenotypes of relapse have been described; in tropical climes relapses occur at short intervals (3-4 weeks), whilst in temperate climes the intervals from primary infection to first relapse may be longer (circa 9 months) [6,7]. Long latency types were prevalent in Europe, North Africa, the former USSR, Central Asia, the Koreas and North and Central America [8], while short latency *P. vivax* malaria types only are found in East Asia and Oceania. Both types coexist in India [9,10].

During the course of repeated *P. vivax* inoculations, individuals living in an endemic area can amass a liver ‘bank’ of genetically diverse hypnozoites [11]. Thus a relapse (i.e. a hypnozoite-derived blood-stage infection) may be caused by parasites that not only differ genetically but also derive from different mosquitoes [11-18]. Standard antimalarials such as chloroquine that are recommended for the treatment of vivax malaria act on blood-stage parasites but they have no effect on the hypnozoites. The only generally available drug that kills hypnozoites and therefore provides ‘radical cure’ is the 8-aminoquinoline primaquine. Although primaquine is generally recommended, it is not used widely because of the risks of iatrogenic haemolysis in patients with glucose-6-phosphate dehydrogenase (G6PD) deficiency [19]. In the absence of radical cure, relapses comprise a substantial proportion of all vivax malaria episodes [6], with Southeast Asia and Oceania having the highest incidence of relapse, and the greatest contribution of relapse to the overall incidence of illness [3,4,20].

Genotyping has proved useful for distinguishing recrudescence and reinfection in the assessment of antimalarial drug efficacy in falciparum malaria, but is not established in the support of efficacy assessments in vivax malaria. Recurrences of *P. vivax* malaria can be caused by recrudescence, reinfection or relapse. The relapses arise from hypnozoites which are derived from the incident infection, or from a previously acquired infection. In order to understand and interpret genotyping outputs, detailed consideration of the biology of malaria infection is required. *Plasmodium* spp. are dioecious parasites, capable of both self and cross-fertilization [21]. During blood-stage infections a sub-population of asexual parasites differentiate into gametocytes, the sexual forms of the parasite [21,22]. Gametocytes ingested by a feeding anopheline vector mosquito undergo gametogenesis, fertilization and consequent sexual recombination [21]. Depending on the number of genetically distinct gametocytes in the blood meal and whether they self or cross-fertilize, the resulting progeny that emerge will exhibit various relationships, described in supplementary Figure 7. These relationships may range from a mixture of unrelated ‘strangers’ to complete clonality.

Finding genotypes compatible with *P. vivax* parasites that are clones or siblings in relation to one another across infections provides strong evidence of either recrudescence or relapse. However, finding genetically unrelated *P. vivax* parasites in a comparison of the acute and recurrent infections is compatible with both relapse and reinfection. Genetic data alone therefore cannot resolve the true ‘state’ (i.e. reinfection, relapse, or recrudescence) of all recurrences since not all genetic data are informative (absence of evidence of relatedness is not evidence of absence of relapse) [23]. In part because relapse is only partially identifiable genetically, previous vivax genotyping work has aimed to distinguish new infection or relapse of heterologous hypnozoites versus recrudescence or relapse of homologous hypnozoites (e.g. [8, 24-26]), where heterologous and homologous are synonymous with an-isogenic and isogenic respectively (see Appendix). While homologous and heterologous are useful genetic descriptors, in therapeutic assessments the primary concern is whether the recurrent infection is a relapse, recrudescence or reinfection.

A major complementary source of information regarding relapse, recrudescence and reinfection is the time-to-recurrence (i.e. interval since treatment of the initial episode). Both short and long latency *P. vivax* relapses exhibit strong periodicity [6]. In the case of recrudescent infections their emergence will be related to parasite biomass at start of treatment, drug pharmacokinetics, host immunity and local resistance patterns. Simple intrahost pharmacodynamic models of malaria argue that relapse will preempt recrudescence when resistance is low grade [6]. Reinfection rates will be either constant over time or seasonal. Time-to-event modeling therefore uses valuable independent information which complements the genetic data.

The inability to distinguish between relapses and reinfections has several consequences for the understanding of the epidemiology of *P. vivax* malaria and the assessment of treatment efficacy. First, routinely collected case data cannot be used to estimate the force of infection, which is an important measure of transmission intensity and a key metric for disease surveillance. In particular, the relationship between force of infection and the number of *P. vivax* cases is non-linear because of relapses [27]. Second, the inability to distinguish between relapses and reinfections means that in endemic populations, estimates of the efficacy of radical curative drugs (i.e. primaquine, the only radical cure currently available, and tafenoquine, approved in some countries but yet to be deployed in endemic areas) in populations at continued risk of infection will be biased downwards by the background transmission rate. Finally, the inability to distinguish between relapse and recrudescence (recurrence of the blood-stage infection as result of blood-stage treatment failure) means that it is difficult to estimate the curative efficacy of a blood-stage treatment in endemic areas [28]. This contrasts with *Plasmodium falciparum* malaria where standardised genotyping is used effectively to distinguish recrudescence from reinfection, allowing for the adjustment of efficacy estimates in populations at continued risk of reinfection [29-31].

In this study, we combined genetic information with longitudinal epidemiological data from two clinical trials of antimalarials in patients infected with *P. vivax*. This was used to test a novel methodological framework that can discriminate between the causes of recurring *P. vivax* infections. We estimated the probabilities of each of these causes: relapse, reinfection and recrudescence, using a generative Bayesian model that combined data on the time-to-recurrence, the drug treatment (taking into account the varying pharmacokinetics and pharmacodynamics of antimalarial drugs), and parasite genetics as characterized by genotyping 3 to 9 highly polymorphic microsatellite markers (repeat length polymorphisms). This analysis pooled data from over 1200 patients collected in two randomized controlled trials on the Thailand-Myanmar border, comparing schizontocidal and hypnozoiticidal drugs for the treatment of *P. vivax* malaria, both with one year follow-up of patients. We showed that by combining *P. vivax* genotyping and time-to-event information relapses could be distinguished reliably from reinfections in this low transmission setting.

## Results

In the VHX study patients of all ages were randomised to receive either artesunate, chloroquine or chloroquine and primaquine [4]. In the BPD study patients were randomised first to receive either chloroquine or dihydroartemisinin-piperaquine and second to receive either 14 days of primaquine 0.5mg/kg/day or 7 days of primaquine 1mg/kg/day [32]. All doses were supervised in both studies. These individual patient-level pooled data contained a total of 2708 time-to-recurrence intervals for 1299 patients (more than 1000 patient-years of combined follow-up time including right censored intervals). The median number of observed recurrences was 2 following both artesunate monotherapy and chloroquine monotherapy, and 0 following high-dose primaquine plus partner drug. In patients not receiving radical cure, 1309 recurrences were observed over 330 patient years of total follow-up; in those receiving radical cure, 130 recurrences were observed over 675 patient years of total follow-up. High-dose primaquine treatment was therefore associated with a 95% decrease in the number of observed recurrences.

### Dynamics of recurrent infections

Because of the periodicity of relapse, and the rarity of recrudescence emerging more than two months after administration of efficacious treatments, the interval between successive episodes of *P. vivax* was shown to be highly informative. We applied a Bayesian population time-to-event mixture model to the 2708 time-to-recurrence intervals. Mean posterior probabilities of the recurrence states (recrudescence, relapse, and reinfection) generated under this model varied over 3 orders of magnitude as a function of time since last episode and treatment drug (Fig 1). Following high-dose primaquine and partner drug, a recurrence in the first few months had a high likelihood of being a relapse, and subsequent recurring infections were almost entirely caused by reinfections (Fig 1).

**Fig 1.**
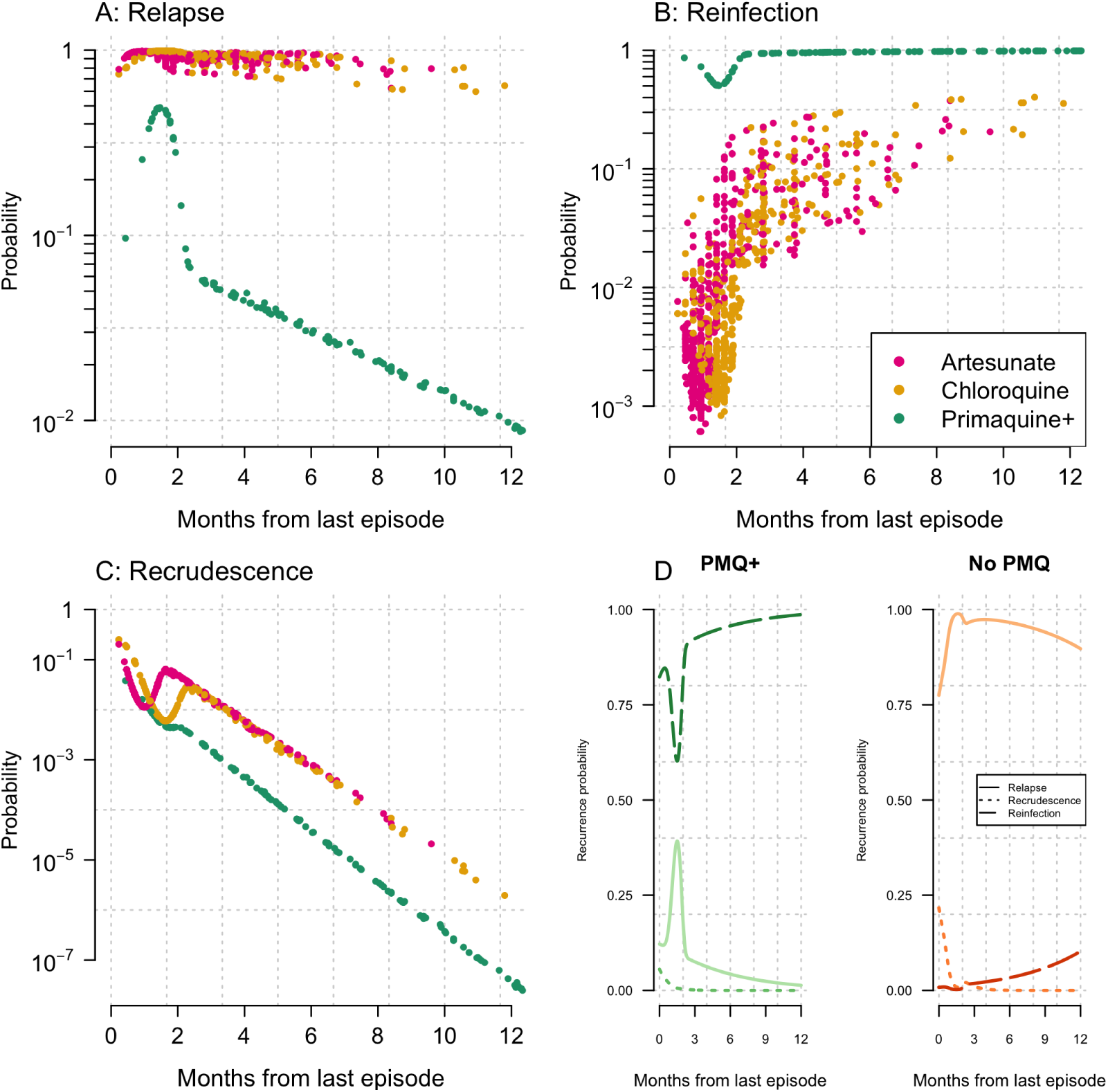
Dynamics of recurrent infections. Estimates of the probabilities on the log_10_ scale of relapse (A), reinfection (B) and recrudescence (C) for all observed recurrences (n=1441) are shown as a function of interval since last episode of vivax malaria (dots). Colours correspond to the treatment used in the previous episode, where Primaquine+ refers to primaquine with partner drug. D: population mean posterior probabilities (normal scale) for the three recurrences types as a function of time-to-recurrence following high-dose primaquine with partner drug (PMQ+) and no primaquine but a slow acting blood-stage drug such as chloroquine (No PMQ).

The posterior uncertainty intervals for the individual probabilities of relapse for each recurrence are shown in Fig 2. For approximately 75% of the recurrences observed after treatment without high-dose primaquine, the posterior distributions were extremely narrow with the probabilities of relapse very close to 1 (Fig 2, top left). The remaining 25% all had relapse posterior probabilities greater than 0.3 but with wide credible intervals. For the recurrences observed after high-dose primaquine, approximately 15% had mean probabilities greater than 0.1 of being relapses and the remaining 85% had mean probabilities less than 0.1 of being relapses (Fig 2, top right panel). In both cases, the time-to-recurrence is correlated with the posterior uncertainty (Fig 2, bottom panels). The least uncertainty was observed around the peak expected timing of relapse following treatment. This is dependent on whether a slowly eliminated blood-stage drug was administered (chloroquine or dihydroartemisin-piperaquine) or a rapidly eliminated (artesunate monotherapy), irrespective of whether high-dose primaquine was added (Fig 2, D).

**Fig 2.**
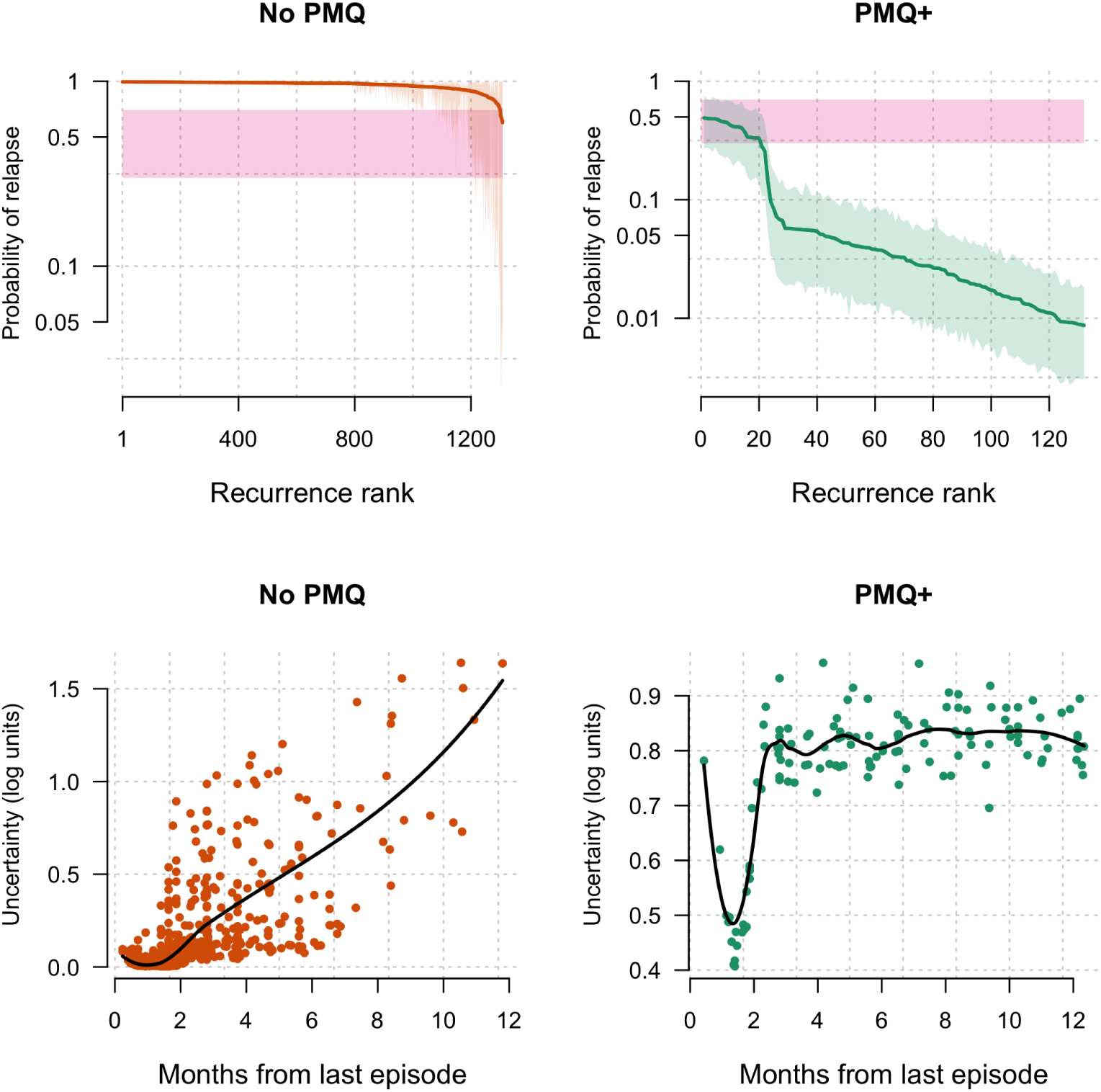
Individual probabilities of relapse based from time-to-event pooled analysis of the VHX and BPD studies. Top panels: per-episode mean probability of relapse along with 95% credible intervals (shaded grey) for 1309 recurrences after no primaquine (left) and for 130 recurrences after high-dose primaquine (right). The recurrences are ranked by their mean probabilities of relapse state. A zone of ‘uncertainty’ (same as in Fig 3) is highlighted in pink. The upper and lower bounds are arbitrary. Bottom panels: the relationship between time since last episode and treatment, and the uncertainty of the posterior estimates (width of the 95% credible interval on the log_10_ scale) after no primaquine (no PMQ, left) and after high-dose primaquine plus partner drug (PMQ+, right). The black lines represent fitted LOESS curves to highlight trends.

The mean estimates of the recurrence states averaged over all observed recurrences in the pooled time-to-event analysis are given in Table 1. Overall, after supervised high-dose primaquine in this epidemiological context, the model estimated that > 90% of the recurrences are reinfections, as compared to less than 4% when radical cure is not given. There was little evidence of recrudescence for all treatments considered. These results are consistent with previous modelling results from the same area [33].

**Table 1.** Mean (95% credible intervals) posterior probability estimates of overall recurrence states in the different treatment arms. All estimates have been rounded up to one decimal place. *Blood stage drug: chloroquine or dihydroartemisinin-piperaquine.

| Treatment | Artesunate<br>monotherapy | Chloroquine<br>monotherapy | Primaquine<br>with blood stage<br>drug* |
| --- | --- | --- | --- |
| Patient years follow-up | 161 | 169 | 675 |
| Recurrences | 722 | 587 | 130 |
| Reinfection (%) | 2.1 (1.4-3) | 3.1 (1.8-3.9) | 90.5 (88.1-92.9) |
| Relapse (%) | 95.6 (94.2-96.7) | 95.5 (94.3-96.7) | 9.4 (6.9-11.8) |
| Recrudescence (%) | 2.3 (1.4-3.5) | 1.7 (1.1-2.6) | 0.2 (0.1-0.3) |

This pooled analysis confirms the high periodicity of relapse for the frequent relapse phenotype present in Southeast Asia [6]. Some previous statistical models of vivax relapse have assumed a constant rate of awaking hypnozoites [27,33-35]. Our pooled analysis takes into account post treatment prophylaxis and shows that the pattern of relapsing infections does not fit a simple constant rate model. The time-to-event model estimates that 60% of relapses can be explained by a periodic Weibull distribution, with the remaining 40% explained by a constant rate exponential distribution. Such a mixture of distributions would fit an ‘activation’ hypothesis, as given by [6], where relapses could generally emerge at a constant rate with a superimposed illness induced feedback (i.e. fever itself activates hypnozoites), thus increasing the probability of activation and creating periodicity.

### Combining time-to-event and genetic information

In order to obtain more informed individual-level estimates of each recurrence state we incorporated information from highly polymorphic microsatellite marker data. We developed a genetic model to describe the relationships between parasites across infections within individuals, using the posterior probabilities of recurrence states from the time-to-event model as the prior. The estimates from the genetic model alone (uniform prior) are shown in supplementary Fig 8. A total of 710 episodes (of which 494 were recurrent) were genotyped from 217 individuals (BPD: n=167 in 80 individuals; VHX: n=543 in 137 individuals). We estimated recurrence state probabilities for 486 of the 494 recurrent episodes (enrollment data were missing for one from BPD, while computational complexity under the genetic model prevented analysis of 7 from VHX). The complexities of infection (COI) in isolates from recurrent vivax episodes were significantly lower than for isolates from enrollment episodes (mean COI of 1.3 versus 1.5, p=0.001). There was no significant difference between the COI of recurrences following high-dose primaquine and those following no primaquine treatment.

Using the genetic model of parasite relatedness between infections, we estimated that in individuals who did not receive high-dose primaquine, nearly all (99.3%, 80% credible interval, CI: 96.8-99.9) of the typed recurrences were relapses (n=365). In contrast, for individuals who were given high-dose primaquine, only 14.3% (80% CI: 12.3-16.7) of recurrences were estimated to be relapses (n=121). The estimates for recrudescence were very low: 0.3% (0.1-0.6) and 0% (0-0.3) for no primaquine and primaquine groups, respectively. Overall, the vast majority of recurrent episodes for which we had genetic data had low uncertainty in the probabilities of their recurrence state (the uncertainty is shown by the vertical lines in Fig 3). We note that trial summaries based on probabilities of the individuals who did not receive high-dose primaquine (all were in the VHX study and the majority received chloroquine monotherapy) are biased by selective genotyping of individuals with the highest number of recurrences. These are presumably the individuals with the largest number of liver hypnozoites and so will not be representative of the general enrollment population. In particular, there are likely competing risks between relapsing infections and reinfections. Relapsing infections will usually reach patency before reinfections, and if they occur simultaneously then the genetic signature could point to relapse. Thus frequent relapse likely hides reinfection. Under the genetic model, we made the simplifying assumption that recurrence states are mutually exclusive.

**Fig 3.**
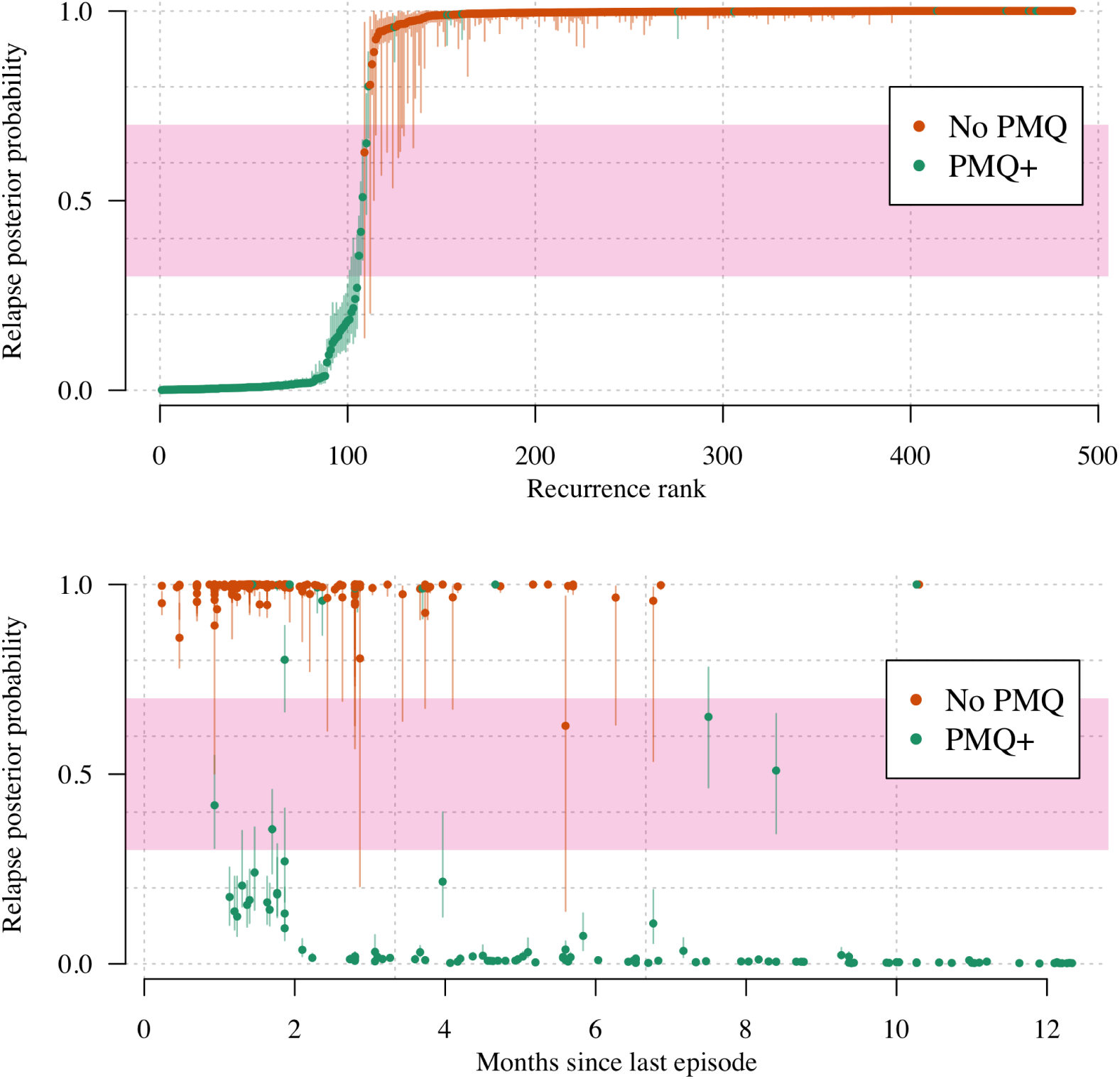
Posterior probabilities of relapse for 486 genotyped *P. vivax* recurrences. The top panel shows the recurrences ranked by their probabilities of relapse state coloured by treatment drug (orange: blood stage treatment only; green: high-dose primaquine plus partner drug). Credible intervals are shown by the vertical lines. The bottom panel shows the same posterior probabilities as a function of the time since the last episode of *P. vivax* with the same color coding. The ‘uncertainty zone’ (same as in Fig 2 used to classify recurrences in Fig 4) is shown by the pink zone (the upper and lower bounds are arbitrary). PMQ: primaquine; PMQ+: primaquine plus partner drug.

Of particular interest are two recurrences in two separate patients which were classified with high certainty as relapses. Both occurred after a 300 day infection free interval (Fig 3, bottom panel). One individual had received high-dose primaquine and the other had not (Fig 4).

**Fig 4.**
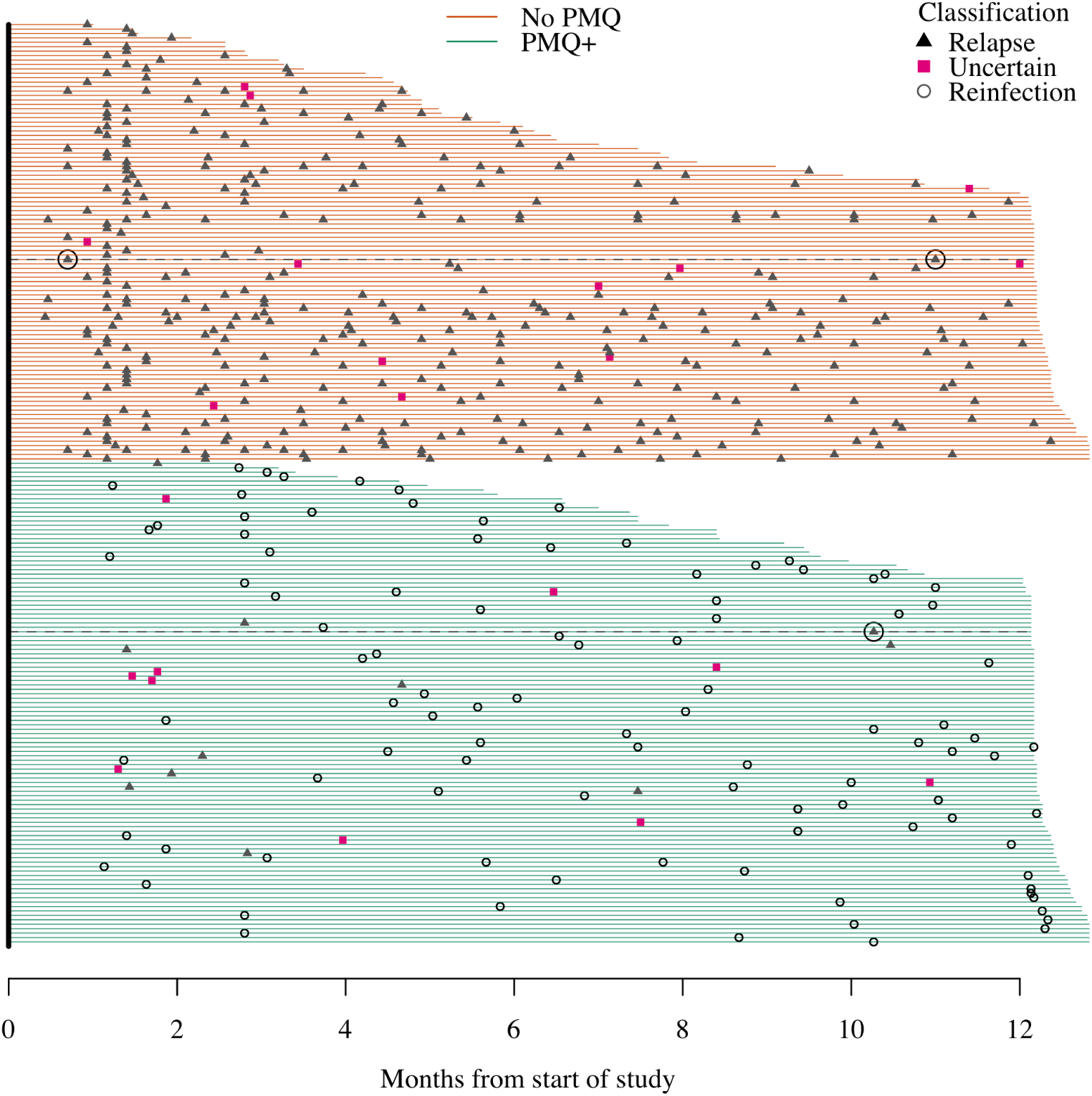
Classification of 486 *P. vivax* recurrences as a function of time since the start of study. Each line represents one individual (n=208). Duration of active follow-up is shown by the span of the horizontal lines (green: high-dose primaquine given; red: no primaquine given). Recurrences classified as relapses are triangles, reinfections are hollow circles, and uncertain classification are pink squares since they fall in the pink zone of uncertainty in Fig 3. The delayed relapses (circa 300 days after treatment) are circled for clarity with their follow-up duration shown by a black dotted line. PMQ: primaquine; PMQ+: primaquine plus partner drug.

We estimated the false-positive discovery rate of the genetic model of recurrence by calculating the probabilities of the relapse state when comparing isolates from episodes in separate individuals. This resulted in 90194 pairwise comparisons with data truly generated under the null distribution: i.e. the parasites in the pairwise comparisons are known to derive from different people and thus cannot be relapses, and genetic data are drawn from the true population distribution. This gave an estimated false positive rate (defined as the probability of relapse greater than the upper bound of an arbitrarily defined ‘uncertainty zone’ from 0.3 to 0.7) of 2.2%. Not only does this highlight the discriminatory power of our panel of 9 microsatellites, but it also highlights the considerable population diversity within a small geographic location with low seasonal transmission.

### Radical cure efficacy of primaquine on the Thailand-Myanmar border

We estimated the reinfection adjusted failure rate of high-dose (total dose of 7mg/kg) supervised primaquine to be 2.6% (80% CI: 2.0-3.5), using the genetic and time-to-event model. This was derived from the final model estimate that 15.8% of recurrences following high-dose primaquine in the BPD study were relapses. If we consider the background transmission to be constant over the 4 years studied, this translates into 99% of all relapses were averted, decreasing the relapse rate from 3.32 per year to 0.03 per year (Table 1). This estimate of number of relapses averted is a function of the individual hypnozoite loads and therefore of the background transmission. These estimates of the radical curative efficacy of high-dose primaquine used all available data from the 655 individuals enrolled in the BPD study, which had requisite genetic data for all but 5 of 92 recurrences. This reinfection adjusted estimate is a considerable reduction from the original reinfection unadjusted estimate of the failure rate at 12% (80% CI: 10-14) [32].

To investigate these results further, we assessed the contribution of individual patient drug exposures by examining the relationship between the day 7 trough concentrations of carboxy-primaquine (the slowly eliminated inactive metabolite of primaquine), and treatment failure (defined as a probability of relapse or recrudescence greater than 0.5), adjusted for primaquine regimen administered (either 14 daily doses of 0.5mg/kg or 7 daily doses of 1mg/kg). A statistically significant trend was observed, but this was driven by a few outliers, defined as episodes in which the plasma carboxy-primaquine trough concentrations were more than 3 standard deviations below the mean (supplementary Fig 9). Concentrations this far below expected values are likely to reflect incomplete drug absorption resulting from protocol deviations (e.g. non adherence, vomiting). After adjusting for these outliers, there was no statistically significant relationship between drug exposure and radical cure failure. This result suggests that it is possible to discriminate between drug failures due to biological mechanisms (e.g. high hypnozoite load, cytochrome P450 2D6 polymorphisms, intrinsic drug resistance, etc.) and drug failures because of vomiting the medication or non-adherence. This is important for correct estimation of drug efficacy. Given the very low failure rate of supervised high-dose primaquine (estimated at 2.6%), only very large pooled patient data analyses would have the necessary power to confirm or refute this conjecture.

### Number of microsatellite markers for reliable assessment of the unknown recurrence states

For microsatellite genotyping in future studies, it is important to determine the minimum number of markers necessary for a reliable assessment of the unknown recurrence states. We simulated data assuming microsatellite markers were independent with effective cardinality of 13 (this is the average effective cardinality in our panel of nine microsatellites, see Methods and Fig 6). To emphasize clearly the information content for a given number of markers in various complexities of infections, we used a uniform distribution over the recurrence states (i.e. recrudesence, reinfection and relapse each have prior probability of ⅓). We simulated paired infections (one primary episode followed by a single recurrence) under three scenarios: the recurrence is either an exact clone, a sibling, or a stranger with respect to the primary infection.

**Fig 5.**
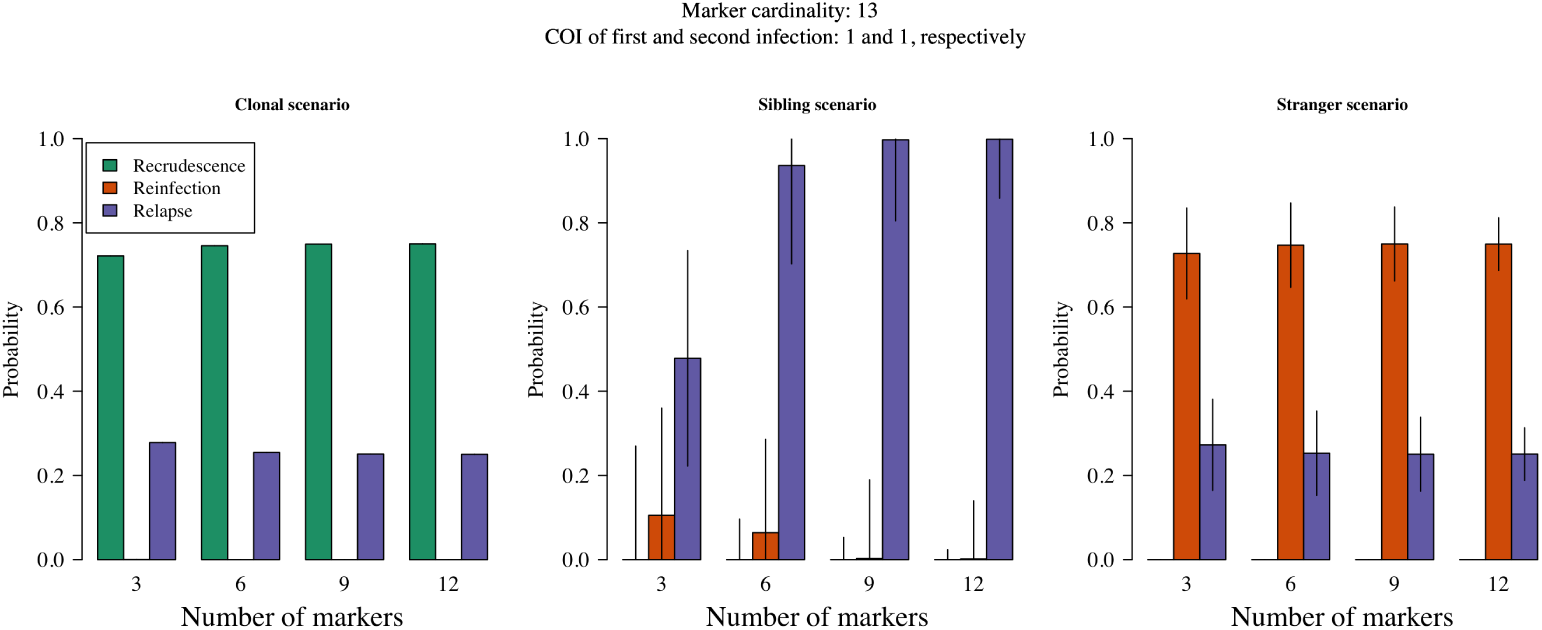
Posterior probabilities of recurrence states as the number of microsatellites typed is increased. Each plot corresponds to a different relationship simulation scenario. Coloured bars show the median posterior probabilities with error bars extending ± one standard deviation. The effective cardinality of each marker was set equal to 13 in these simulations. The COIs in the first and second infections were 1 and 1 respectively. The prior probability of each recurrence state was ⅓. As the number of microsatellites genotyped is increased, the following are expected. 1) Under the *clone scenario*, the probability of recrudescence should converge a probability greater than the prior and the complement of that of relapse; while the probability of reinfection should converge to 0. 2) Under the *sibling scenario*, the probabilities should converge to 1 for relapse and 0 otherwise. 3) Under the *strange scenario*, the probability of reinfection should converge to a probability greater than the prior and the complement of that of relapse; while the probability of recrudescence should converge to 0.

**Fig 6.**
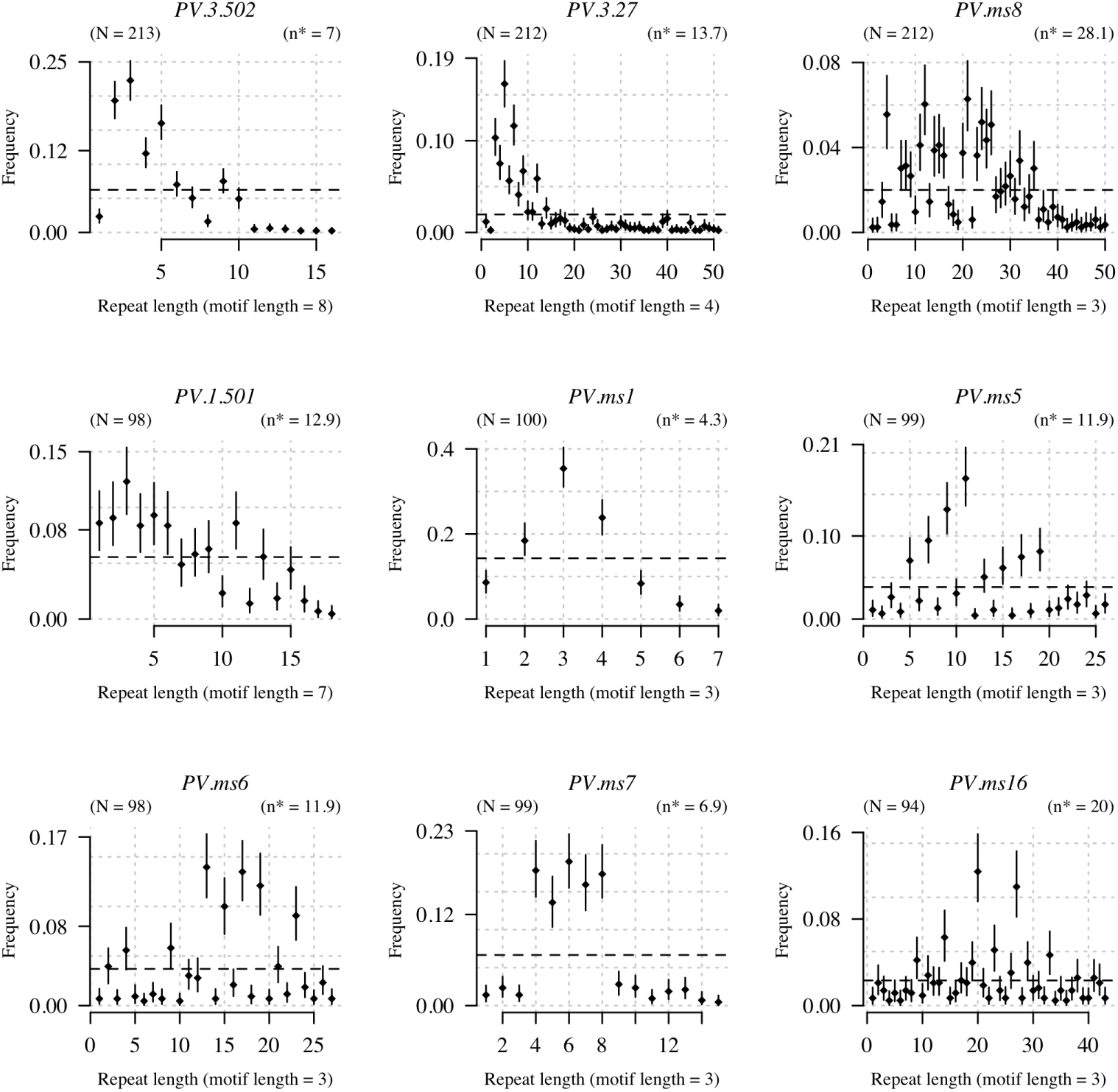
Allele frequencies for the 9 microsatellites estimated using enrolment episodes from 216 individuals (79 in BPD and 137 in VHX)). The mean frequency estimate is shown by the black circles and 95% credible intervals are shown by the vertical lines. The dotted horizontal line shows the discrete uniform distribution over alleles for comparison. The number of isolates typed for each microsatellite is denoted by *N*. The top row shows the microsatellites which were typed most frequently *(PV.3.502, PV.3.27, PV.ms8*). The range of feasible alleles is given by the maximum observed repeat length in all the data per microsatellite combined, and all repeats are given a pseudo-observation of weight *w* = 1. The estimated effective cardinality of each marker is denoted by *n*^*^.

Fig 5 shows the posterior probabilities of the recurrence states for each of these three scenarios as a function of the numbers of markers typed. When the simulated data had clonal or ‘stranger’ relationships, six or more markers sufficed to recover expected probabilities. When the simulated data had sibling relationships (exclusive evidence of relapse) nine markers or more were needed to obtain a median posterior probability of relapse close to one. These simple simulations suggest that reliable posterior estimates of the unknown recurrence state can be obtained with approximately 9 markers of effective cardinality equal to 13.

## Discussion

Distinguishing recrudescence, relapse and reinfection in recurrent vivax malaria is necessary for optimum planning of malaria control and elimination interventions as each recurrence state has a different implication for case management. To date, there are no available host or parasite biomarkers of the different recurrence states. Our approach is to our knowledge the first principled attempt to use treatment information along with time-to-event and genetic data to estimate individual probabilities of the possible recurrence states. The most important operationally relevant result from this analysis is a re-evaluation of the estimated radical cure failure rate following high-dose primaquine. We re-estimated the true failure rate in the BPD study to be 2.6% as compared to the published reinfection unadjusted estimate of 12% [32]. The radical curative efficacy of primaquine is not a fixed property - even when reinfections are discounted, it depends on hypnozoite burden and thus the background level of transmission [6, 34] - so these results also probably reflect recent improvements in malaria control in the area [36]. Our analysis reinforces the value of high-dose primaquine radical cure in this setting.

We note that the estimated false positive discovery rate for relapse was extremely low (2.2%) when the genetic model alone was tested on isolates from separate patients (null data). This guards against underestimating the efficacy of high-dose primaquine. Additionally, the estimated number of relapses in non-primaquine treated patients was very high (exceeds 98%). This means that the false negative rate for relapse identification in non-primaquine treated patients cannot be more than around 2% and tentatively suggests a similarly low rate for relapses in patients who do receive primaquine, which guards against overestimating the efficacy of high-dose primaquine.

Much remains unknown regarding the biology of relapse and the mode of action of the 8-aminoquinoline radical curative drugs. No good *in vitro* hypnozoite model is yet available. Under a simple *in silico* model of relapse fit to the data from trials on the Thailand-Myanmar border, where *P. vivax* exhibits the short-latency phenotype [6,7], we recovered an approximate 60:40 split between early-periodic relapse and constant rate relapse. Whether this truly captures the biology of relapse activation or whether a more complex system operates is uncertain. We also observed two late recurrences (300 days since last episode) with high probability (and low estimate uncertainty) of relapse. Our results thus suggest that short-latency hypnozoites can remain dormant in the liver for up to year, confirming previous reports with presumably similar *P. vivax* phenotypes (the Chesson strain) [5,13]. These late relapsing hypnozoites likely awaken via a different mechanism to that of the highly periodic ‘long-latency’ *P. vivax* [6]; the most parsimonious explanation would be that they awake at ‘random’ (constant rate of relapse).

Various statistical and mathematical models of *P. vivax* exist, ranging from models of within host dynamics to global geostatistical descriptions of prevalence [7,27,33-35,37-54]. However, only two publications to our knowledge have formally considered recurrence states on an individual basis [37,55]. Ross *et al*. modelled data from a cohort study in Papua New Guinea (an area of substantially higher transmission than the Thailand-Myanmar border, or indeed almost any vivax endemic area) to estimate *P. vivax* seasonality and incidence [55]. The model is based on the presence-absence patterns of alleles at two polyallelic markers, each modelled separately, allowing for various mechanisms that beget unobserved alleles, but neither reinfection with the same genotype nor recombination. As such, relapses caused by genetically distinct hypnozoites (siblings or ‘strangers’) are unaccounted for. White *et al*. combined genetic (from a single polyallelic locus) and time-to-event data using a statistical model of parasite clone acquisition and clearance using samples from cohorts in Papua New Guinea and Thailand [37]. The majority of recurrent episodes were asymptomatic and not treated, and the main target of inference was the duration of asymptomatic infection. Individual probabilities of relapse were estimated but not fully identifiable. Data on a single locus did not discriminate between a single blood-stage infection and multiple successive relapses.

This is the first attempt to apply a principled probabilistic model framework of relapse, recrudescence and reinfection in recurrent vivax malaria using both time-to-event and multi-locus genetic data on multiple episodes of malaria in the same individual. It allows complementary information from different data types to be quantified systematically. However, strong assumptions were necessary (a comprehensive list can be found in the Methods) and the genetic model has limitations. The main limitations are poor ability to infer a recrudescent state, and computational complexity. In general, correct classification of recrudescent infections is difficult as at low levels of resistance they will reach patency at similar times as relapsing infections and the genetic signature is the same as a homologous (i.e. isogenic) hypnozoite. Our genetic model is brittle with respect to recrudescence as we do not account for imperfect detection or genotyping errors. Given there is very little evidence of recrudescence in this area from other indicators (i.e. slowing of parasite clearance rates and rising recurrence rates supported by reduced in-vitro susceptibility), this is unlikely to impact on our results, but it would necessitate modification before application to data from a region where *P. vivax* antimalarial resistance is suspected. The computational complexity of the model is exponential as a function of the number of recurrences and the complexities of the episodes. Its scope is therefore limited to data from a maximum of three episodes (a primary episode and two recurrences) in which the maximum cumulative COI is 6, with approximately 3-6 heteroallelic genotype calls per episode. The majority of *P. vivax* endemic areas are consistent with the low transmission settings studied here on the Thailand-Myanmar border [1], where over 99% of the single recurrences fit these stipulations (the median COI in both VHX and BPD is 1). However, the method would require modification before application to data from high transmission settings such as Papua New Guinea, where the estimated complexity of a single infection is often greater than 6 [37,55].

Transmission intensity correlates positively with both genetic complexity and diversity [56]. Our model implies a trade-off: decreasing genetic complexity decreases computational complexity, but decreasing genetic diversity decreases the resolving power of the genetic markers. *P. vivax* has high levels of genetic diversity even in low transmission settings, probably as a result of ‘heterologous’ genotype activation [6,57-59]. Nevertheless there is a transmission intensity ‘sweet spot’ for inference whereby genetic complexity is sufficiently low and diversity sufficiently high.

A simple simulation study estimated that genotyping 9 highly polymorphic microsatellites was sufficient for reliable inference of recurrence states. There are no commensurable models of vivax recurrence with which to compare this result. It agrees roughly with estimates from parentage and sibship studies [60,61], where both parent-sibling and sibling relationships have an expected relatedness of 0.5 (supplementary Figure 7). Highly polyallelic microsatellite length poymorphisms have long been used for relatedness inference due to their per-locus resolving power, but in suitably equipped laboratories they are being superseded by whole genome sequencing (WGS) data [62-64]. WGS data provide greater sensitivity to resolve partially related parasites, while providing additional information on population genetics and possibly drug resistance markers [26,65], but are relatively costly. Moreover, the information they provide may exceed that required for recurrence state inference, which is primarily concerned with a subset of relationships: clones, siblings and strangers. With this in view, there has been recent interest in using micro-haplotypes for recurrence state inference. Micro-haplotypes combine the resolving power of highly polyallelic microsatellites with high-throughput ease, thereby addressing economic, technical and statistical concerns [66,67].

Models based on the time-to-event fit well generally and can provide probabilistic estimates of hidden recurrence states. However, they necessitate large training samples (the pooled analysis trial used over 2700 time intervals recorded in over 1200 individuals) and do not use important information regarding relatedness as captured by genetic data. On the other hand, genetic data alone can generate estimates of relapse, recrudescence and reinfection, but they are largely uninformative when pairs of infections are unrelated. Alone, each model is useful but sub-optimal. In combination they provide more informed probabilistic estimates. In this work we combined the two models informally, using the posterior of the time-to-event model as a discrete prior in the genetic model. Using this approach we determined that the radical curative efficacy of supervised high-dose primaquine is considerably higher than previously thought in the epidemiological setting of frequent relapse vivax malaria on the Thailand-Myanmar border.

## Conclusion

Time-to-recurrence combined with highly polymorphic microsatellite genotyping data provide valuable complimentary information for the inference of relapse, recrudescence and reinfection states of recurrent vivax infections.

## Methods

### Clinical procedures

This section summarizes the clinical procedures followed in the Vivax History (VHX) trial [4] and the Best Primaquine Dose (BPD) trial [32]. Both trials were conducted by the Shoklo Malaria Research Unit in clinics along the Thailand-Myanmar border in northwestern Thailand. This is an area with low seasonal malaria transmission. The patient populations comprised of migrant workers and displaced persons of Burman and Karen ethnicity [68]. During the time these studies were conducted, primaquine radical cure treatment was not routinely given to patients in this area.

### Ethical Approval

The BPD study was approved by both the Mahidol University Faculty of Tropical Medicine Ethics Committee (MUTM 2011-043, TMEC 11-008) and the Oxford Tropical Research Ethics Committee (OXTREC 17-11) and was registered at ClinicalTrials.gov number NCT01640574. The VHX study was given ethical approval by the Mahidol University Faculty of Tropical Medicine Ethics Committee (MUTM 2010-006) and the Oxford Tropical Research Ethics Committee (OXTREC 04-10) and was registered at ClinicalTrials.gov trial number NCT01074905.

### Vivax History trial (VHX)

This randomized controlled trial was conducted between May 2010 and October 2012. 644 patients older than 6 months and weighing more than 7kg with microscopy confirmed uncomplicated *P. vivax* mono-infection were randomized to receive artesunate (2 mg/kg/day for 5 days), chloroquine (25 mg base/kg divided over 3 days: 10 mg/kg, 10 mg/kg, 5 mg/kg) or chloroquine plus primaquine (0.5 mg base/kg/day for 14 days). G6PD abnormal patients (as determined by the fluorescent spot test) were randomized only to the artesunate and chloroquine monotherapy groups. The aim was to measure chloroquine efficacy and characterise the ‘natural’ history of *P. vivax* recurrence in order to assess the risks and benefits of radical cure.

Subjects were followed daily for supervised drug treatment. Follow-up continued weekly for eight weeks and then every four weeks for a total of one year. Patients with microscopy confirmed *P. vivax* infections were retreated with the same study drug as in the original allocation. Patients in the artesunate monotherapy or chloroquine monotherapy groups who experienced more than 9 recurrences were given radical curative treatment with the standard primaquine regimen (0.5 mg base/kg/day for 14 days).

### Best Primaquine Dose trial (BPD)

Between February 2012 and July 2014, 680 patients older than 6 months were enrolled in a four-way randomized controlled trial simultaneously comparing two regimens of primaquine (0.5 mg/kg/day for 14 days or 1 mg/kg/day for 7 days) combined with one of two blood-stage treatments: chloroquine (25 mg base/kg) or dihydroartemisinin-piperaquine (dihydroartemisinin 7mg/kg and piperaquine 55mg/kg). The aim was to characterize the efficacy and tolerability of a shorter course (7 days rather than 14) of higher dose primaquine, with a second randomization to chloroquine or dihydroartemisinin-piperaquine. All doses were supervised.

The inclusion and exclusion criteria for this study were the same as for the VHX study, except for the following: patients were excluded if they were G6PD deficient by the fluorescent spot test, had a haematocrit less than 25%, had received a blood transfusion within 3 months, or could not comply with the study requirements.

Follow-up visits occurred on weeks 2 and 4, and then every 4 weeks for a total of one year. Any recurrent *P. vivax* infections detected by microscopy (same criteria as for VHX) were treated with a standard regimen of chloroquine (25 mg base/kg over 3 days) and primaquine (0.5 mg base/kg/day for 14 days).

In both studies, recurrent episodes were detected actively at the scheduled visits by microscopy (lower limit of detection is circa 20 parasites per μL). Patients were encouraged to come to the clinics between scheduled visits when unwell and so some recurrences were detected passively (less than 5%).

### Microsatellite genotyping

Whole blood for complete blood count was collected by venipuncture in a 2mL EDTA tube. Remaining whole blood was frozen at −80C. *Plasmodium vivax* genomic DNA was extracted from 1 mL of venous blood using automated DNA extraction system QIAsymphony SP (QIAGEN, Germany) and QIAsymphony DSP DNA mini kit (QIAGEN, Germany) according to the manufacturer’s instructions. In order to compare primary infections and recurrences genotypic patterns, we genotyped initially using three polymorphic microsatellite loci that provided very clean amplification: no stutter peaks, and on the basis of amplification (i.e. reliability of PCR amplification at the low parasite densities usually found in recurrent infections). These loci were PV 3.27, PV 3.502, and PV ms8. PCR amplification was performed following previously described protocols [12,69]. The genotypes of pre- and post-treatment samples were subsequently assigned a crude classification: ‘related’ versus ‘different’. ‘Related’ genotypes were defined as identical alleles observed at at least two loci out of the three typed. If all alleles at all loci were different, or identical alleles were only observed at one loci, the samples were classified as ‘different’. If the paired samples were classified as ‘related’, six more microsatellite markers were genotyped (PV 1.501, PV ms1, PV ms5, PV ms 6, PV ms7, and PV ms16).

For allele calling on the microsatellites, the lengths of the PCR products were measured in comparison to internal size standards (Genescan 500 LIZ) on an ABI 3100 Genetic analyzer (PE Applied Biosystems), using GENESCAN and GENOTYPER software (Applied Biosystems) to measure allele lengths and to quantify peak heights. Multiple alleles were called when there were multiple peaks per locus and where minor peaks were > 33% of the height of the predominant allele. We included negative control samples (human DNA or no template) in each amplification run. A subset of the samples (n = 10) were analyzed in triplicate to confirm the consistency of the results obtained. All pairs of primers were tested for specificity by use of genomic DNA from *P. falciparum* or humans.

### Statistical models

All model code and statistical data analysis were written in R (version 3.4.3). The genetic model uses the R package *igraph* [70] to manipulate relatedness graphs. Time-to-event models were written in *rstan* based on the *stan* probabilistic programming language [71,72]. Logistic and poisson mixed effects regression models were fitted using the package *lme4*. R code along with deindentified microsatellite and time-to-event datasets can be found on github at github.com/jwatowatson/RecurrentVivax.

### Time-to-event model of vivax recurrence

For recurrent *P. vivax* infections in the VHX and BPD studies, we developed and compared two Bayesian mixed-effects mixture models describing the time-to-event data conditional on the treatment drug administered. Notation was chosen so as to be consistent with the mathematical notation for the genetic model (next section). For each individual (denoted by the subscript *n* ∈ *1*‥*N*), we record the time intervals (in days) between successive *P. vivax* episodes (the enrollment episode is denoted episode 0). The last time interval is right censored at the end of follow-up. The models assume early dropout is not a confounding factor for drug efficacy. For the *n*^*th*^ individual, data concerning the time interval *t* (the time between episode *t —* 1 and episode t) is of the form 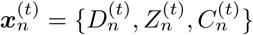, where 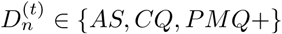 is the drug combination used to treated episode *t* — 1 (AS: artesunate monotherapy; CQ: chloroquine monotherapy; PMQ+: primaquine plus blood stage treatment), 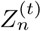 is the time interval in days, and 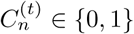, where 1 corresponds to a right censored observation (i.e. follow-up ended before the next recurrence was observed) and 0 corresponds to an observed recurrent infection. In general, let 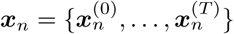, denote all available time-to-event data for the *n*^*th*^ individual. Few recurrences (n=8) occurred in the first 8 weeks for patients randomized to the dihydroartemisinin-piperaquine arms of the BPD trial, so we modelled the post-prophylactic period of piperaquine as identical to that of chloroquine (i.e. PMQ+ includes both chloroquine and dihydroartemisinin-piperaquine as blood stage treatments). In reality the elimination profiles and intrinsic activities are slightly different, with piperaquine providing slightly longer asexual stage suppression than chloroquine.

Time-to-recurrence is modeled as a mixture of four distributions, with mixture weights depending on treatment of the previous episode. The mixture distributions correspond to the different recurrence states. The four mixtures are: *reinfection* given by an exponential distribution; *early/periodic* relapse given by a Weibull distribution with treatment drug dependent parameters; *late/random* relapse given by an exponential distribution; *recrudescence* given by a Weibull distribution with treatment drug dependent parameters.

Model 1 assumes 100% efficacy of high-dose primaquine with only reinfection possible. Model 2 does not assume 100% efficacy of high-dose primaquine and specifies different mixing proportions for the reinfection component in the non primaquine and primaquine groups, *p*_*n*_*(AS)* = *p*_*n*_*(CQ)* and *p*_*n*_*(PMQ+)*, respectively. The mixing proportion between late and early relapse within the relapse component is the same across primaquine and non-primaquine groups. The likelihood for model 2 is given as:

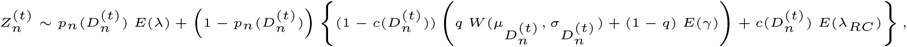

where *p*_*n*_(·) ∈ (0,1) is the individual and drug-specific mixture probability of reinfection (we set the prior to reflect our belief that *p*_*n*_*(AS)* = *p*_*n*_*(CQ) > p*_*n*_*(PMQ+))* and c(·) ∈ (0,1) is the nested drug-specific mixture probability of recrudescence.

The likelihood for model 1 is the same except that *p*_*n*_*(PMQ+)* = 1 (only reinfection possible). λ is the rate of reinfection; *E*(·) denotes the exponential distribution. *E(*λ_*RC*_) is an exponential distribution with rate parameter λ_*RC*_ (assumed drug independent) modelling the time to recrudescence; the parameters 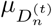 and shape parameter 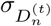 determine the drug-specific mean and variance of the time to relapse, with *μ* _*CQ*_ = *μ* _*PMQ+*_ and *σ* _*CQ*_ = *σ*_*PMQ+*_. *q* is the doubly nested mixing proportion between ‘periodic’ (first component) and ‘random’ (second component) relapses. This is a fixed proportion across all individuals. The ‘random’ relapses are parameterized by the rate constant γ. The individual marginal probability of reinfection is given by 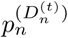; the individual marginal probability of recrudescence is given by 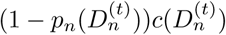; the individual marginal probability of relapse is given by 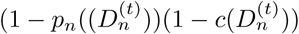.

We used informative prior distributions (supplementary Fig 10) to ensure identifiability of the mixture components. Information content in the data, over and above that specified in the prior, was visually examined using prior-to-posterior plots. The prior-to-posterior plot for model 2 is shown in supplementary Figure 10.

**Fig 8.**
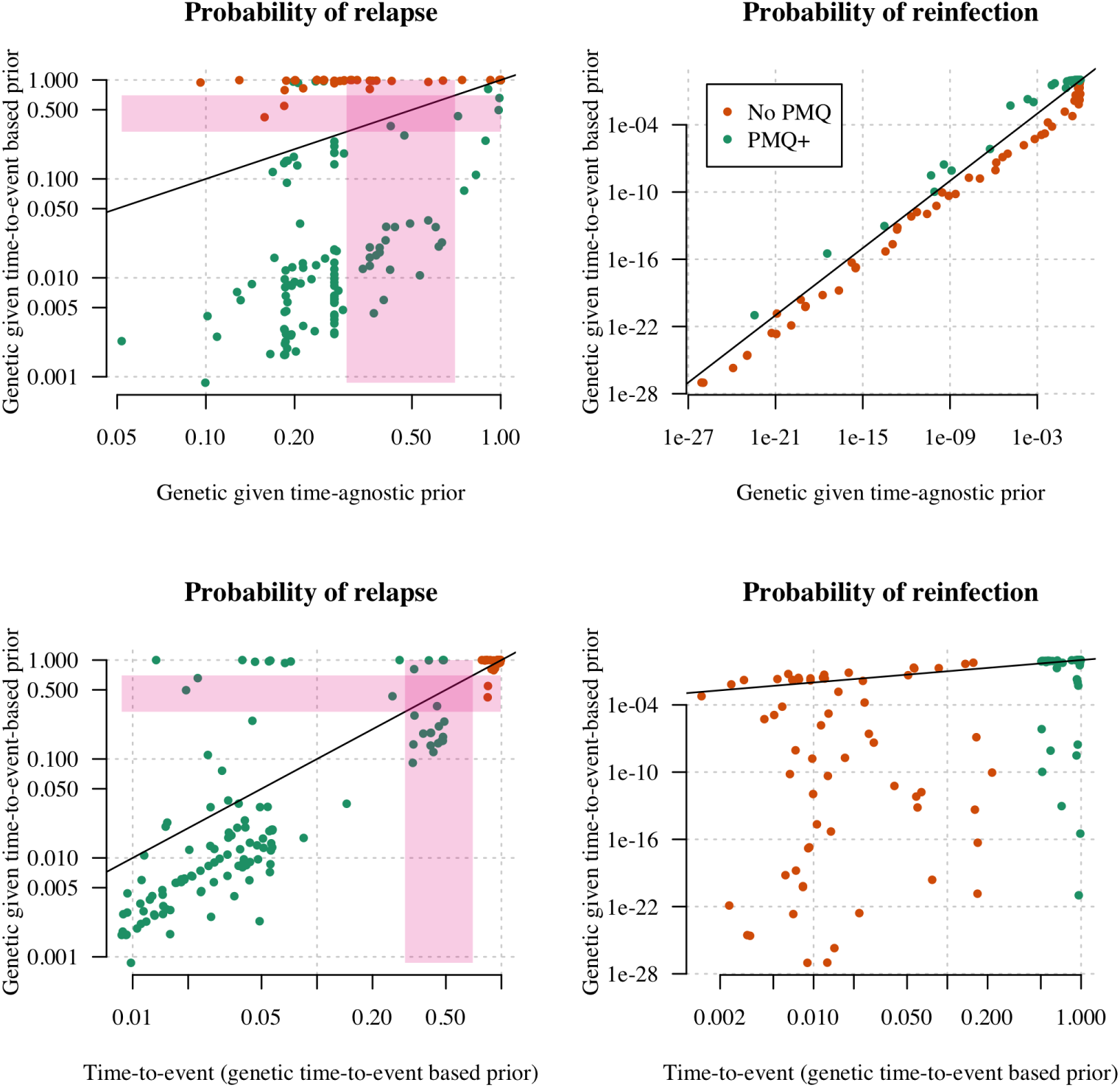
Effect of incorporating both time-to-event and genetic data on probabilities of relapse and reinfection based on either data type alone. The top two plots show the effect of incorporating time-to-event data, relative to genetic data alone, on probabilities of relapse (left) and reinfection (right). The bottom two plots show the effect of incorporating genetic data, relative to time-to-event data alone, on probabilities of relapse (left) and reinfection (right). The diagonal line marks the line of equality. Pink bands mark the arbitrarily defined relapse classification ‘uncertainty zones’. Points in top left and bottom right ‘certainty zones’ (non-pink rectangles) mark recurrences whose classification changes from not relapse to relapse and vice versa, respectively, upon incorporation of data. Upon incorporation of time-to-event data some ‘No PMQ’ and to a lesser extent some ‘PMQ+’ treated recurrences are classified as relapse where they were not previously, while three ‘PMQ+’ treated recurrences are classified as not being recurrences where they were previously (top left plot). Upon incorporation of genetic data, some ‘PMQ+’ treated episodes are classified as relapses where they were not previously.

**Fig 9.**
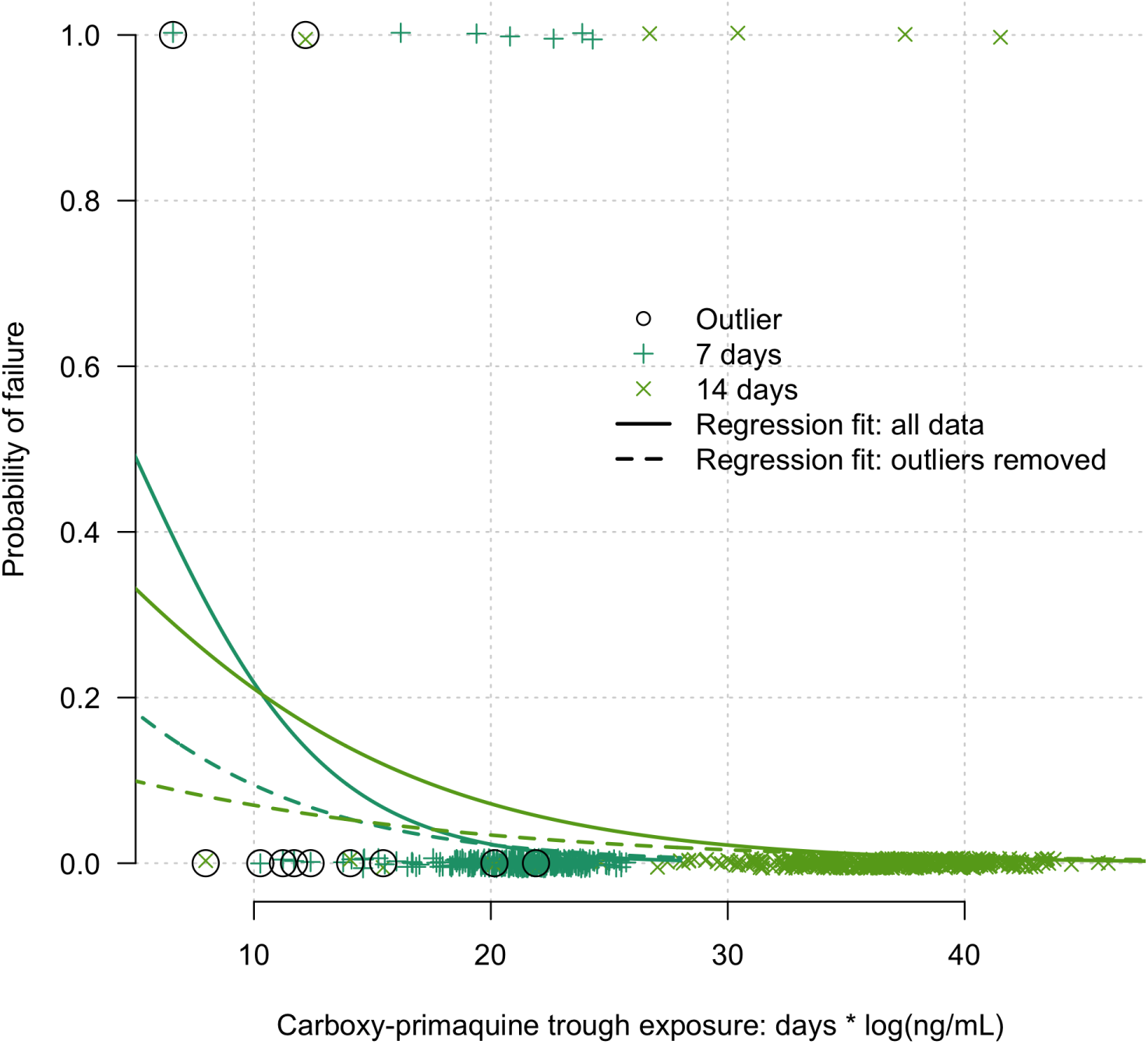
Estimated relationship between carboxy-primaquine exposure and primaquine failure. The proxy exposure to carboxy-primaquine is defined (lower bound) as the number of days of primaquine administration multiplied by the log trough concentration observed on day 7. The fitted trends using all the data are significant; they are shown by the thick lines. After removal of outliers (circles: defined as those episodes whose carboxy-primaquine trough concentrations were more than 3 standard deviations from the mean), the fitted trends (non-significant) are shown by the dashed lines.

**Fig 10.**
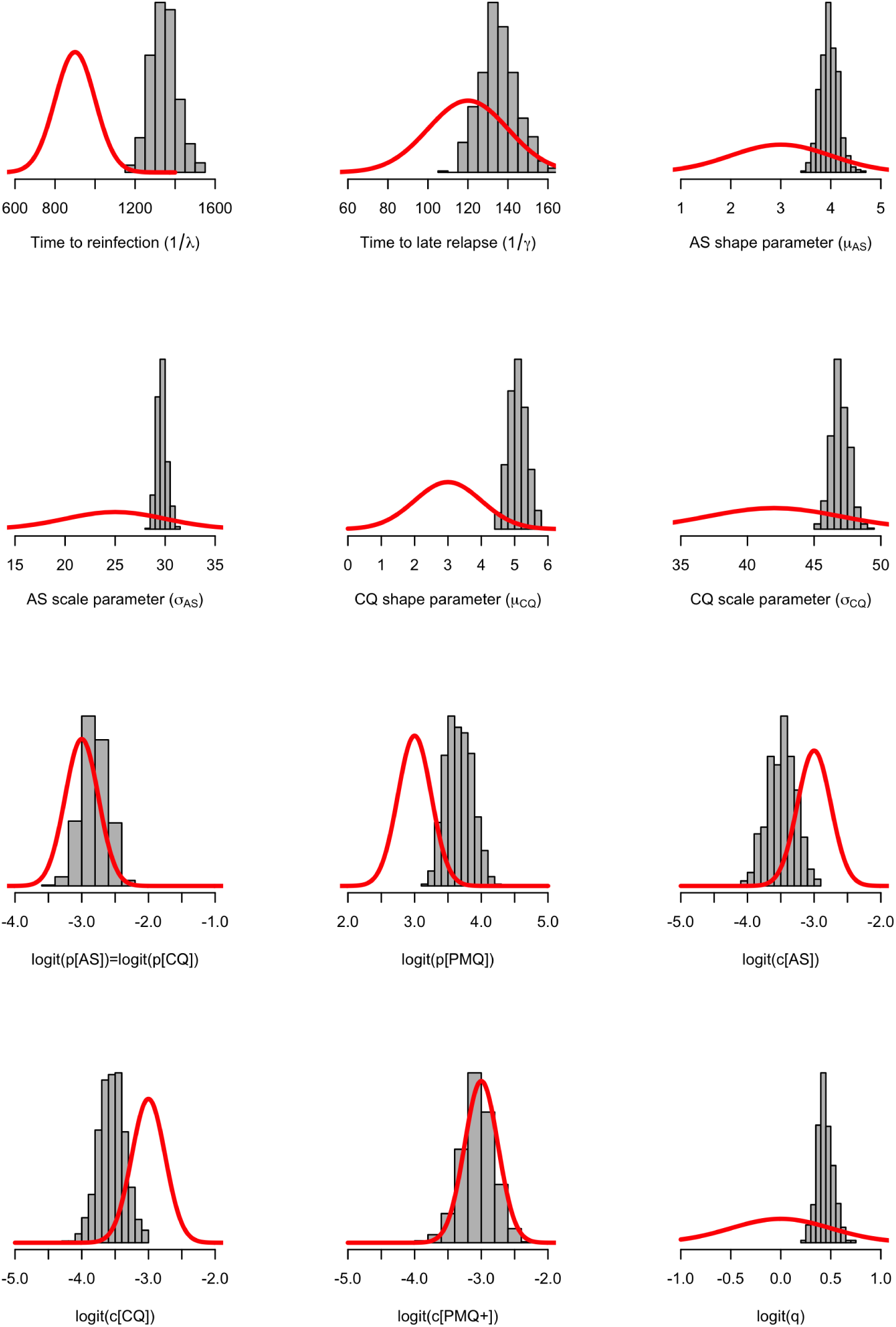
Information contained in time-to-event data. Going from prior distributions (red thick lines) to posterior distributions (normalised histograms) for the population level model parameters. For each prior and posterior, the horizontal axis is density.

All time-to-event models were coded and implemented in *stan* [72] and run using R version 3.4.3. The complete models can be found on github at github.com/jwatowatson/RecurrentVivax/Timing_Model.

The stan models output (i) Monte Carlo posterior distributions for all model parameters; (ii) posterior estimates of recurrence states for each time interval 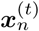; (iii) log likelihood estimates of each posterior draw. For each model, we ran 8 chains with 10^6^ iterations, thining per 4000 iterations and discarding half for burn-in. Convergence of MCMC chains was assessed using traceplots assessing mixing and agreement of the six independent chains. These can be found in the github repository.

Neither of the models incorporate seasonality, which is an important factor in *P. vivax* transmission on the Thailand-Myanmar border. A seasonal model is in development and will be the subject of future work.

### Determination of population allele frequencies

Allele frequencies are critical for assessing the probability of identity by chance between isolates under the genetic model (described below). Given that the two clinical studies (VHX and BPD) were carried out in the same clinics and populations at a similar point in time (both were completed within a 4 year period), to estimate allele frequencies we combined all microsatellite data from the enrollment episodes in the two studies. Specifically, this included 216 enrollment episodes, with 137 individuals from the VHX trial and 79 from the BPD trial. We excluded recurrent episodes since repeat relapses are liable to introduce intra-individual correlation.

There are several ways to compute allele frequencies from enrollment data. A simple approach is to use monoclonal data. However, some alleles are only seen in polyclonal infections. In addition, monoclonal-derived frequencies would almost certainly introduce bias resulting from disproportionately low representation of rare alleles across monoclonal infections [73]. A statistically rigorous treatment would jointly model frequencies, polyclonality and hidden recurrence states, but is prohibitively complex. We therefore took a two step approach: first we generated frequency estimates and then, conditional on these estimates, we modelled relapse allowing for polyclonality (genetic model described below).

To compute allele frequencies we used a multinomial likelihood and Dirichlet prior with weight *ω* = 1 (equivalent to *ω* pseudo observations per allele) both with dimension equal to the maximum repeat length seen across all the data, thereby interpolating unobserved repeat lengths less than the maximum observed. Conjugacy implies the posterior distribution over allele frequencies is a Dirichlet distribution with parameter vector equal to *ω* plus the vector of allele counts (counted as 1 per episode observed). The mean posterior estimate was used as a point estimate for the allele frequencies. A Monte Carlo approximation of an 80% credible interval (extending from the 10th to 90th percentile) was constructed using 1000 random draws around the point estimates (Fig 6).

For each microsatellite genotyped, we calculated its ‘effective cardinality’ n*, defined as the theoretical number of alleles (non-integer) which under a uniform distribution would give rise to the same probability of identity by chance. The estimated values of n* are shown for each microsatellite in Fig 6, with values varying from 4 *(PV.ms1*) to 28 *(PV.ms8*).

To lessen the probability that two alleles are identical by chance, some studies discard common alleles (e.g. [55]). In this study, we account for identity by chance using a model of relatedness based on identity by descent (IBD).

### Genetic model of relapse

In this section we describe the genetic model that outputs the probability that the *t*^*th*^ recurrence of the *n*^*th*^ individual is a recrudescence, relapse or reinfection, given a vector of prior probabilities of recrudescence, relapse or reinfection, and genetic data on a small number of polyallelic microsatellite markers. In our joint model of time-to-event and genetic data, the prior vector in the genetic model is derived from the posterior distribution over recurrence states given the time-to-event data. Comprehensive mathematical details of the model can be found in the Appendix.

To infer the probability of relapse, we construct a statistical model that exploits evidence of relatedness expected within a single mosquito inoculum following recombination (supplementary Fig 7). The only relationships between sporozoites that are exclusive to a single mosquito are those between meiotic clones and siblings (Table 2). In other words, relapse can be caused by many types of sporozoites, including those which are ‘strangers’ in relation to one another, but, in the absence of recrudescence, only meiotic clones and meiotic siblings provide conclusive evidence of relapse (Table 2). Genetic data cannot discern clones and siblings that are meiotic or not. We thus use a coarse definition of clones and siblings in this model, ignoring whether the relationship is meiotic. We use the term ‘stranger’ to refer to all parasites whose shared ancestry dates back beyond the most recent mosquito inoculation. Consequently, the term clone refers to first-generation clones only. We assume zero probability of reinfection with first-generation clones or siblings, since the probability of a mosquito feeding on the same human host consecutively is low. Importantly, we do allow ‘strangers’ that are genetically identical by chance. This captures some clones resulting from clonal expansion (i.e. multiple generations of selfing). Parent-offspring pairs are considered siblings under our model since they have an expected relatedness equal to 0.5 in the absence of inbreeding. We ignore half siblings whose expected relatedness is 0.25 in the absence of inbreeding. However, we do allow half-siblings in the sense that we allow two or more alleles per locus within infections assumed to contain only siblings (i.e. we allow for collections of siblings that together share more than two parents). The main model assumptions (including those mentioned above, also summarised in Table 2) are as follows.

**Fig 7.**
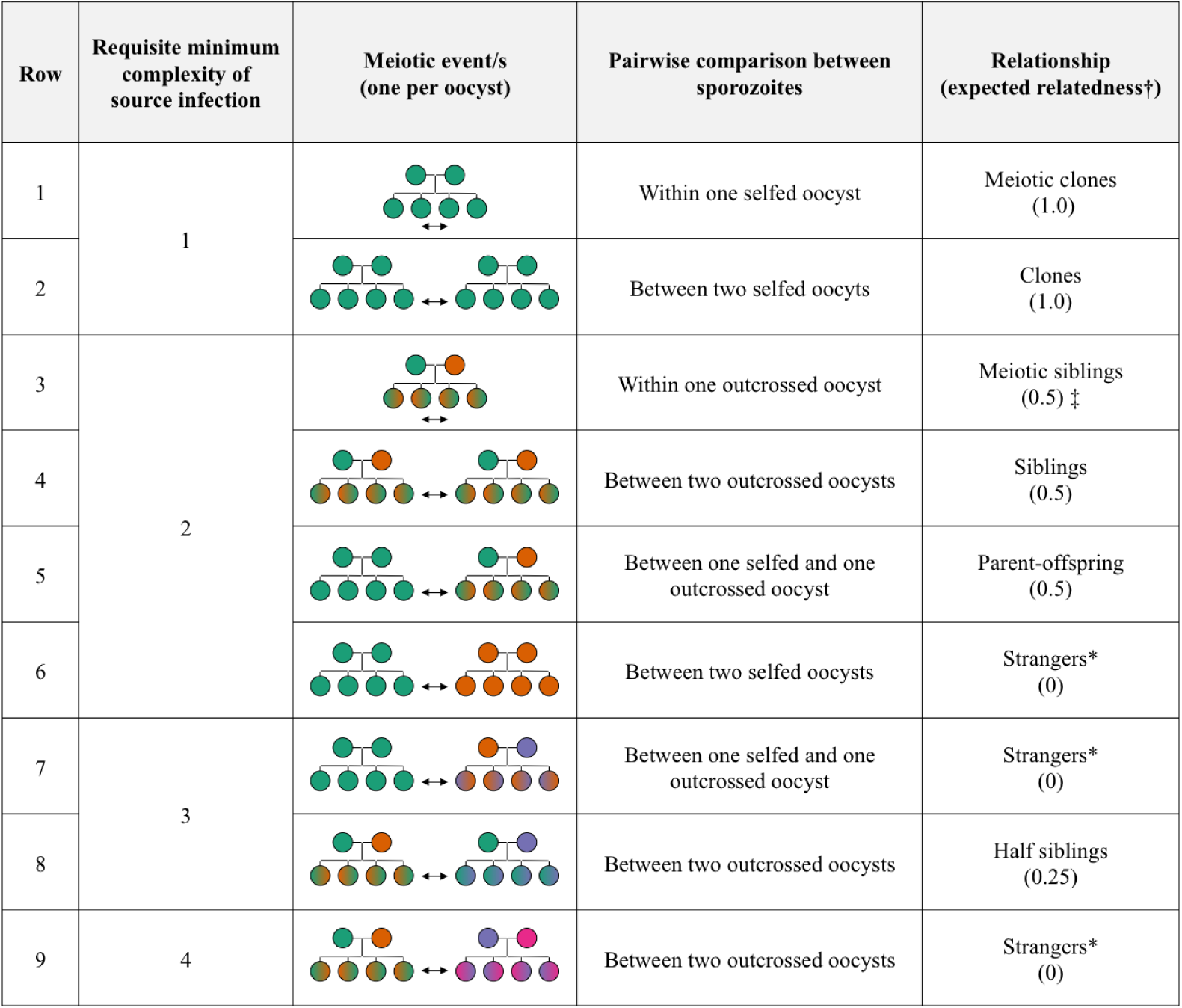
Possible meiotic events and subsequent relationships between sporozoites within a single mosquito. This is an adaptation of Fig 2 of [82]. Possible meiotic events (of which there can be one or more within a mosquito, but only one per zygote resulting in one oocyst) are represented by trees with two root nodes representing parental gametes and four leaf nodes representing recombinant offspring that subsequently replicate by mitosis producing thousands of sporozoites within the oocyst. Different colours represent different genotypes. Hybrid colours represent outcrossed offspring. We show scenarios for a maximum of four genetically distinct parental gametes as this encompasses all possible pairwise relationships. ^†^Expected relatedness values assume a large randomly mating population (i.e. no inbreeding). ^‡^The expected relatedness of meiotic siblings is the mean of a bimodal distribution with modes at 0.3 and 1 (see Appendix and [76,82] for explanation). Strangers* include all parasites whose shared ancestry dates back beyond the mosquito in question.

**Table 2.** Compatibility of parasite relationships *across* blood-stage infections conditional on recrudescence, relapse and reinfection assuming all parasites are detected during patent infection (i.e. underlying truth). Meiotic clones are parasites derived from a selfed oocyst (row 1, supplementary Fig 7), which means they must have been coinoculated. Meiotic siblings are parasites derived from an outcrossed oocyst (row 3, supplementary Fig 7), which also means they must have been coinoculated. Clones, siblings, parent-offspring and half siblings are all parasites that share common gamete genotypes but are from different oocysts (i.e. rows 2, 4, 5, and 8, respectively, supplementary Fig 7). They can occur in a single mosquito or in different mosquitoes (e.g. contemporaneous mosquitoes that share a common human source, or sequential mosquitoes linked to a common human host who acts a source to the first mosquito and sink to the second). Strangers* include all parasites whose shared common ancestry dates back beyond the most recent mosquito (rows 6, 7 and 9, supplementary Fig 7). Under the model, clones^†^ includes both meiotic and not; siblings^‡^ includes parent-offspring and meiotic and not, but excludes half siblings in the sense that they have expected relatedness of 0.25 (in the absence of inbreeding).

|  |  | Recrudescence | Relapse | Reinfection |
| --- | --- | --- | --- | --- |
| Real life | Meiotic clones | Yes | Yes | No |
|  | Meiotic siblings | No | Yes | No |
|  | Clones | No | Yes | Yes but unlikely |
|  | Siblings | No | Yes | Yes but unlikely |
|  | Half siblings | No | Yes | Yes but unlikely |
|  | Parent-offspring | No | Yes | Yes but unlikely |
|  | Strangers* | No | Yes | Yes |
| Model | Clones<sup>†</sup> | Yes | Yes | No |
|  | Siblings<sup>‡</sup> | No | Yes | No |
|  | Strangers* | No | Yes | Yes |

1. In relation to one another, sporozoites within an inoculum are either clones, siblings or ‘strangers’ with respective expected relatedness of 1, 0.5 + α and 0 + α (where α = 0 in the absence of inbreeding, otherwise α ∈ (0, 0.5])
2. Recrudescence has non-zero probability if and only if all parasites are clones of those in the directly preceding infection
3. No genotyping errors (i.e. no microsatellite slippage)
4. No mutation (i.e. all diversity due to standing variation)
5. 100% detection of all parasite clones during patent infection
6. The complexity of a given infection (COI) is equal to the maximum number of distinct alleles seen at any microsatellite within that infection
7. The probability of being reinfected with parasites that share alleles with previously inoculated parasite is equal to that of identity by chance (i.e. the product of the frequencies of the alleles observed).
8. Mutually exclusive recurrence states
9. All microsatellites are independent and neutral

Assumption 1 disregards half-siblings. However, as described above, within infections assumed to contain only siblings, we allow more than two alleles per locus. Half-siblings are possible within an inoculum if both i) a mosquito takes a blood meal from an infection with COI ≥ 3 (supplementary Fig 7), and ii) two or more zygotes derived from the cross fertilization of three or more genetically distinct gametes survive the severe bottleneck of successful oocyst formation [21,74]. Given the severe bottleneck, and the fact that only 30 of the 710 (4%) genotyped episodes in the pooled dataset have COI estimates ≥3, half-siblings are likely to be rare in the data analysed here. If any are present, the likelihood of the observed alleles will be misspecified with expected IBD=0.5 under the model.

Assumption 1 also states that all siblings have the same expected relatedness (0.5 in the absence of inbreeding). In general, this assumption holds for all full siblings, meiotic or not (supplementary Fig 7). However, if we condition on meiotic siblings having one or more genetic differences, the expected relatedness between two meiotic siblings is 0.4 (see Appendix). Within infections with COI≥ 2, we collapse clonal relationships between haploid parasite genotypes. This means that the expected relatedness is very slightly over-specified (by 0.1) if an infection contains meiotic sibling parasites with distinct haploid genotypes. This level of misspecification is likely inconsequential given natural variation introduced by meiosis [75,76]. Moreover, if we average over the probability that the two distinct parasites came from different oocysts (e.g. rows 4, 5, 6, supplementary Fig 7), the extent of misspecification rapidly declines.

To allow for inbreeding under the model, we have included the parameter α ∈ [0, 0.5] as specified in Assumption 1. A recent study found compelling evidence of inbreeding and selfing in oocysts from mosquitoes fed on *P. vivax* blood samples from the Thailand-Myanmar border in 2013 [77]. As a sensitivity analysis, we reran the computations with α= 0.175 (see github notebook) and compared with results for α= 0. The overall results are robust to this change, the only impact being that some primaquine treated recurrences having a lower probability of relapse.

Assumptions 2 to 6 render inference of recrudescence frail under the model. Given there is very little evidence of recrudescence in the time-to-event data, and there is no other evidence to support recrudescences, this likely has little impact on our results. The model would require modification before application to data from a region where *P. vivax* antimalarial resistance is suspected, however. In comparison, inference of relapse under the model is robust to assumptions 3 and 4 because we account for relapses caused by siblings under the model. Conditioning on sibling relatednesss (IBD=0.5) absorbs differences caused by mutation and slippage, especially since within infections assumed to contain only siblings we allow more than two alleles per locus. Assumption 7 is likely to hold except in the rare event that an infected mosquito consecutively feeds on the same human host (thereby transmitting a first-generation clone or recombinant offspring to the same human host from whom the parental parasites were sourced).

Assumption 8 implies that all polyclonal reinfections are generated by co-inoculation. That is, we do not allow coincidental blood-stage parasites from different mosquito inoculations unless they are hynozoite-derived. Both the BPD and VHX trials had active follow-up and all asymptomatic infections were treated so this assumption is likely to hold. It would not hold in the context of passive detection or untreated asymptomatic infections. This assumption also implies that we will miss reinfections in individuals with frequent sibling or clonal relapses.

Neutrality of microsatellites (assumption 9) implies that allele frequencies of hypnozoites are the same as those in the wider population. The microsatellite markers used in this study were designed specifically to meet this assumption [12,69]. Future data types may include non-independent markers. Extension of the relatedness model to capture linkage between non-independent markers is possible (see [76,78,79]).

Informally, the model proceeds by explicitly summing over labelled graphs of relatedness between parasites both within and across infections. Each vertex label represents a parasite haploid genotype, each edge label represents a relationship between a pair of parasite haploid genotypes. To account for multiple haploid genotypes within complex infections while maintaining the most parsimonious representation of the data, the number of vertices per infection within each relatedness graph is set equal to the maximum number of distinct alleles seen at any microsatellite within that infection (i.e. the COI, assumption 6). Conditional on this number of vertices we enumerate all possible ways to phase the microsatellite data (i.e. all possible ways to label vertices with parasite haploid genotypes); enumerate all viable relationships between parasite genotypes (i.e. all possible ways to label edges with relationships), allowing either sibling or stranger edges within infections (clonal edges within infections collapse), and clone, sibling or stranger edges across infections. For each combination of vertex and edge labels, we then compute the probability of the data conditional on the expected relatedness given the edge label and the allele frequencies of the vertex labels. We integrate over states of IBD, thereby taking into account alleles that are identical due to chance. We allow for some background IBD, since serial transmission of related parasites can lead to higher than expected IBD in low transmission areas, using the parameter α (discussed above). Finally, we sum over all relatednesss graphs weighted by their probability given relapse, recrudescence or reinfection.

The probability of a relatedness graph given that the recurrence is a relapse is equal over all viable graphs, since all viable relatedness graphs are compatible with a relapse state (Table 2). The probability of a relatedness graph given the recurrence is a recrudescence is equal over viable graphs that have, for each haploid parasite genotype in the recurrent infection, one or more clonal edges with haploid parasite genotypes in the previous infection, since only clones are compatible with recrudescence under the model (Table 2 and assumption 2). The probability of a relatedness graph given the recurrence is a reinfection is equal over viable graphs that have only stranger edges between infections, since clones and siblings are not compatible with a reinfection state under the model (Table 2 and assumption 7).

For a given recurrence state, we do not weight the probability of each graph by the relative proportion of vertices that are clones, siblings and ‘strangers’ in relation to one another. Doing so is theoretically possible (and could provide a link to a dynamic *P. vivax* transmission model), but is very difficult without prior knowledge of the relative proportions expected given different recurrence states. A full understanding requires joint modelling of the hidden recurrence states, transmission (to capture the expected counts of ‘strangers’ and siblings in co-inoculations, thus treating COI as a random variable) and of the impact of host covariates (e.g. age, treatment history, etc.) on the hypnozoite bank to better understand the observed variance in homologous versus heterologous relapse [12,14-18,24,80].

### Genetic model for more than 2 recurrences

Due to computational complexity, the genetic model of recurrence is limited to the joint analysis of one or two recurrences only (see Discussion). This amounted to n=158 patients in the pooled dataset. For individuals with more than two recurrences (n=54), we estimated all within individual pairwise probabilities of recurrence states between episodes (1178 pairwise comparisons). The results of these pairwise comparisons can be used to construct for each individual an adjacency matrix of recurrence states between episodes. Relapse probabilities are then defined as proportional to the maximum estimated probability of relapse with respect to all preceding episodes. Recrudescence probability is proportional to the probability of recrudescence with respect to the directly preceding episode. For each recurrence, the probability of reinfection is the complement of the probability of relapse plus recrudescence.

### Joint model of relapse

#### Motivation

Time-to-event data often only provide intermediate level evidence for or against relapse and ignore rich signals from genetic data. Alone, genetic data do not suffice to pinpoint relapsing infections as unrelated parasites are found both within and across inocula, and are compatible with relapse or reinfection. However, by combining both sources of information we can use genetic data to update the *a priori* belief of recurrent states based on the time-to-event. In this work we combine the two models informally, using the posterior of the time-to-event model as a discrete prior in the genetic model. It remains to be seen whether a formally joint model of both data types would add value [81].

#### Conditional independence of time-to-event and genetic information

For the *n*^*th*^ individual, let *x*_*n*_ denote all available time-to-event data (as above), *y*_*n*_ denote all available genetic data, and ***R***_*n*_ denote hidden recurrent states. Under the joint model of relapse we assume *x*_*n*_ and *y*_*n*_ are conditionally independent given ***R***_n_:

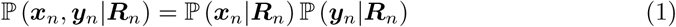

This assumption allows sequential update of the posterior probability of ***R***_*n*_. First given information from the time-of-event data,

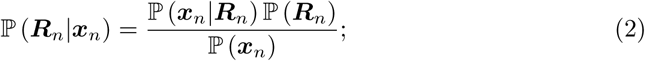

second given information from genotyping (be it microsatellite data or other),

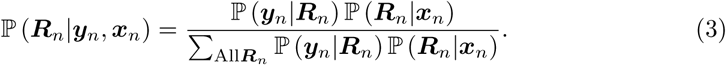

We use Monte Carlo sampling to numerically approximate 𝕡 (***R***_*n*_*|x*_*n*_), Eq 2. We then draw from the numerical approximation uniformly at random (while also drawing from the posterior allele frequency distributions) to recover a numerical approximation of 𝕡 (***R***_*n*_*|x*_n_, *y*_*n*_). Since we explicitly sum over all ***R***_*n*_, for a given allele frequency draw, (Eq 3) can be considered a transformation of the sample approximating 𝕡 (***R***_*n*_|x_*n*_).

The conditional independence assumption can be interpreted as ‘no propensity for relapsing stranger parasites to occur at different time intervals than relapsing parasites that are related’. In reality, this assumption may be incorrect. For example, [6] hypothesises that malarial illness itself activates pre-existent hypnozoites which in long latency *P. vivax* could lead to a preferential activation of genetically unrelated hypnozoites [6]. In long latency vivax, reinfection would thereby trigger activation of previously inoculated parasites (unrelated) and the most recently accumulated hypnozoites would stay dormant for 8-9 months. However, this is highly speculative and thus in our joint model of relapse we assume conditional independence.

### Classification of recurrent episodes

The estimation of the false positive discovery rate of the genetic model and Figure 4 both necessitate the specification of classification boundaries. We arbitrarily chose the interval [0.3,0.7] as the ‘zone of uncertainty’, with probabilities of a recurrence state greater than 0.7 implying a certain classification.

## Acknowledgments

NJW is supported by a Principal Investigator award from the Wellcome Trust. KP is supported by the Royal Golden Jubilee Ph.D. Programme, the Thailand Research Fund (PHD/0032/2556). ART and COB were supported by a NIGMS Maximizing Investigator’s Research Award (MIRA) R35GM124715-02. This project has been funded in part with Federal funds from the National Institute of Allergy and Infectious Diseases, National Institutes of Health, Department of Health and Human Services, under Grant Number U19AI110818 to the Broad Institute (DN).

The content is solely the responsibility of the authors and does not necessarily represent the official views of the funders.

We are grateful to all the patients who took part in these studies and for the study staff who cared for them. A special thanks to Clare Ling, PhD and Pornpimon Wilairisak for managing and keeping in order the large volume of study samples.

**S1 Plot Possible relationships between sporozoites within a single mosquito inoculum**

**S2 Plot Contribution of different data sources to final estimates**

**S3 Plot Estimated relationship between carboxy-primaquine exposure and primaquine failure.**

**S4 Plot Information contained in time-to-event data** Going from prior distributions (red thick lines) to posterior distributions (histograms) for the population level model parameters.

## Appendices

### Terminology: homologous and heterologous

Typically, the term “homologous” is synonymous with genetically identical or near identical (i.e. allowing for genotyping error), whereas “heterologous” is synonymous with genetically different. Both terms are used interchangeably as descriptors both on the level of a parasite haploid genotype (e.g. ‘heterologous hypnozoite’ [12]) and on the level of an infection (e.g. ‘homologous recurrences’ [9]). When referring to an infection, the terms apply to collections of parasites, which may contain different genotypes that are not necessarily resolved. They are likely unresolved because most genotyping methods. For example, if two alleles are detected per locus at two loci in a primary infection (alleles 1 and 2 at locus A; alleles 3 and 4 at locus B) then we do not know how these ‘phase’ to parasite haploid genotypes (do we have A1B3 and A2B4, or A1B4 and A2B3?), and if the recurrence contained the same four alleles, we cannot therefore be sure we have the same parasites (i.e. the primary infection could be A1B3 and A2B4, and the recurrence could be A1B4 and A2B3). Parasites that are siblings, and also the infections that they cause, could be deemed homologous (as in [6]), heterologous (as in [26,28]), or referred to separately (as in [18,65,83]).

### Relatedness between meioitic siblings

As discussed in [82], the average pairwise relatedness between sporozoites that are meiotic siblings is 0.5, despite average pairwise relatedness between haploid meiotic products (hereafter referred to as HMPs) being 0.33. This is because of the massive expansion of the HMPs during sporogeny [21,74]. In short, expansion amounts to there being 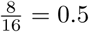 IBD permutations of sporozoites from the mature oocyst, despite 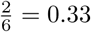 IBD combinations of HMPs from the tetraploid zygote. A more detailed explanation follows.

Fig 11 is a schematic of sexual recombination between genetically distinct malaria parasites based on the review [21]. Different colours denote different genomes. The microgamete is blue, and the macrogamete is red. A single locus is highlighted by a solid circle. In step 1, two genetically distinct haploid gametes (one micro, one macro, represented by blue and red, respectively) come together forming a diploid zygote. Endomeiotic replication follows, resulting in a tetraploid zygote with four presumed HMPs (step 2). The zygote then transforms into a motile ookinete before maturation into an oocyst, where, in a process called sporogony, the four presumed haploid meiotic products replicate by endomitosis producing thousands of genomes that mature into sporozoites (step 3).

**Fig 11.**
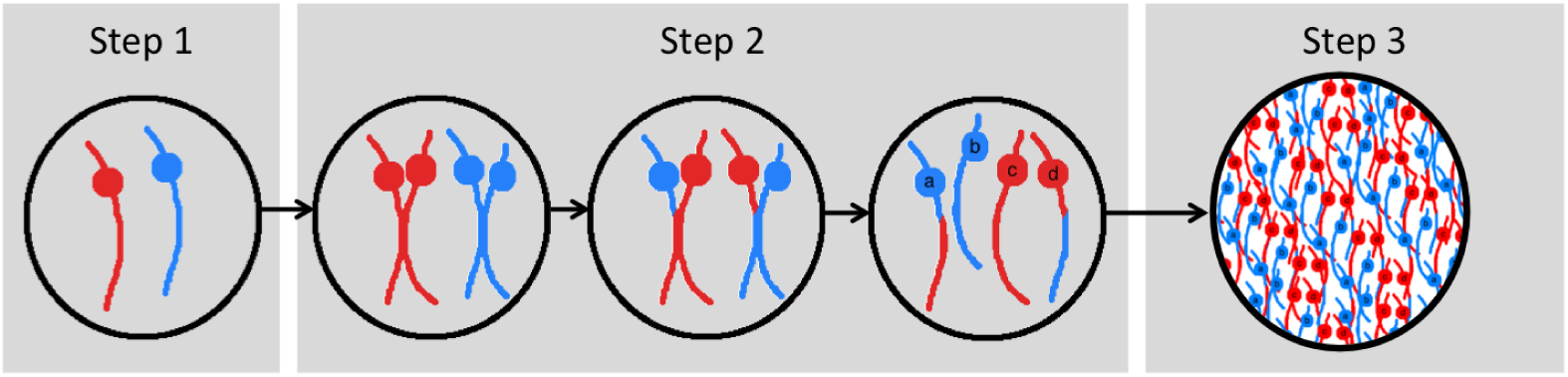
A schematic of sexual recombination between genetically distinct malaria parasites based on a review by [21]. Different colours denote different genomes. A single locus is highlighted by a solid circle. Two haploid gametes (one micro, one macro) fuse forming a diploid zygote (step 1). Endomeiotic replication follows, resulting in a tetraploid zygote with four presumed HMPs, labelled a, b, c and d (step 2). Note that the labels a to c refer to the HMPs in their entirety, not the alleles at the highlighted locus. The zygote then transforms into a motile ookinete, where the second round of meiosis is thought to occur, before maturation of the ookinete into an oocyst. Within the oocyst, the four presumed haploid meiotic products replicate by endomitosis producing thousands of genomes that mature into sporozoites (step 3).

Since there are only four HMPs in the tetraploid zygote (a, b, c and d in Fig 11), there are only 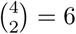 ways to select distinct pairs of HMPs (ab, ac, ad, bc, bd and cd). A pair is identical by descent (IBD=1) at a given locus only if the HMPs share a marker inherited from the same gamete (otherwise IBD=0). At any given locus, 2 HMPs inherit markers from the microgamete (a and b at the highlighted locus in Fig 11), there is thus 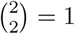 way of choosing two markers derived from the microgametes (ab in this example). Similarly, there is 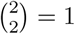 way of selecting two markers derived from the macrogamete (cd in this example). Of the 6 HMP pairs, there are thus two that are identical by descent (IBD=1: ab and cd) and 4 that are not (IBD=0: ac, ad, bc, bd), amounting to an average pairwise relatedness between HMPs of 2/6 = 0.33.

Since within an oocyst there are thousands of sporozoites derived from only 4 HMPs we can select sporozoites derived from the same and different HMPs (i.e. equivalent to sampling HMPs with replacement). Given the four HMP precursors, there are in total 4 x 4 = 16 ways to select sporozoites (aa, ab, ac, ad, ba, bb, bc, bd, ca, cb, cc, cd, da, db, dc, dd). Given two of the four HMP precursors inherit markers from the microgamete (a and b), there are 2 x 2 = 4 ways two select two sporozoites that have markers derived from the microgametes (aa, bb, ab, ba). Similarly there are 2 x 2 = 4 of selecting two markers derived from the macrogamete (cc, dd, cd, dc). Of the 16 sporozoite pairs, 4 + 4 = 8 are thus IBD, amounting to an average pairwise relatedness between sporozoites of 8/16 = 0.5.

If we condition on sporozoites having one or more genetic differences (e.g. exclude repeats aa, bb, cc, dd from numerator and denominator), the average pairwise relatedness between remaining sporozoites is 4/10 = 0.4. This means that the expected relatedness between two genetically distinct sporozoites that are meiotic siblings is 0.4. On the opposite end of the spectrum, if genetically identical micro and macro gametes self-fertilize all combinations and permutations are IBD and the haploid meiotic products are clonal.

### Genetic model: full mathematical description

In this section we provide a full mathematical description of the genetic model. It is broken down into four subsections entitled: *Model overview, Evaluation of the prior, Evaluation of the likelihood* (divided into three parts), *Model implementation*. A running example is provided in boxes. The full set of mathematical notation used in this model is given in Table 3.

**Table 3.**
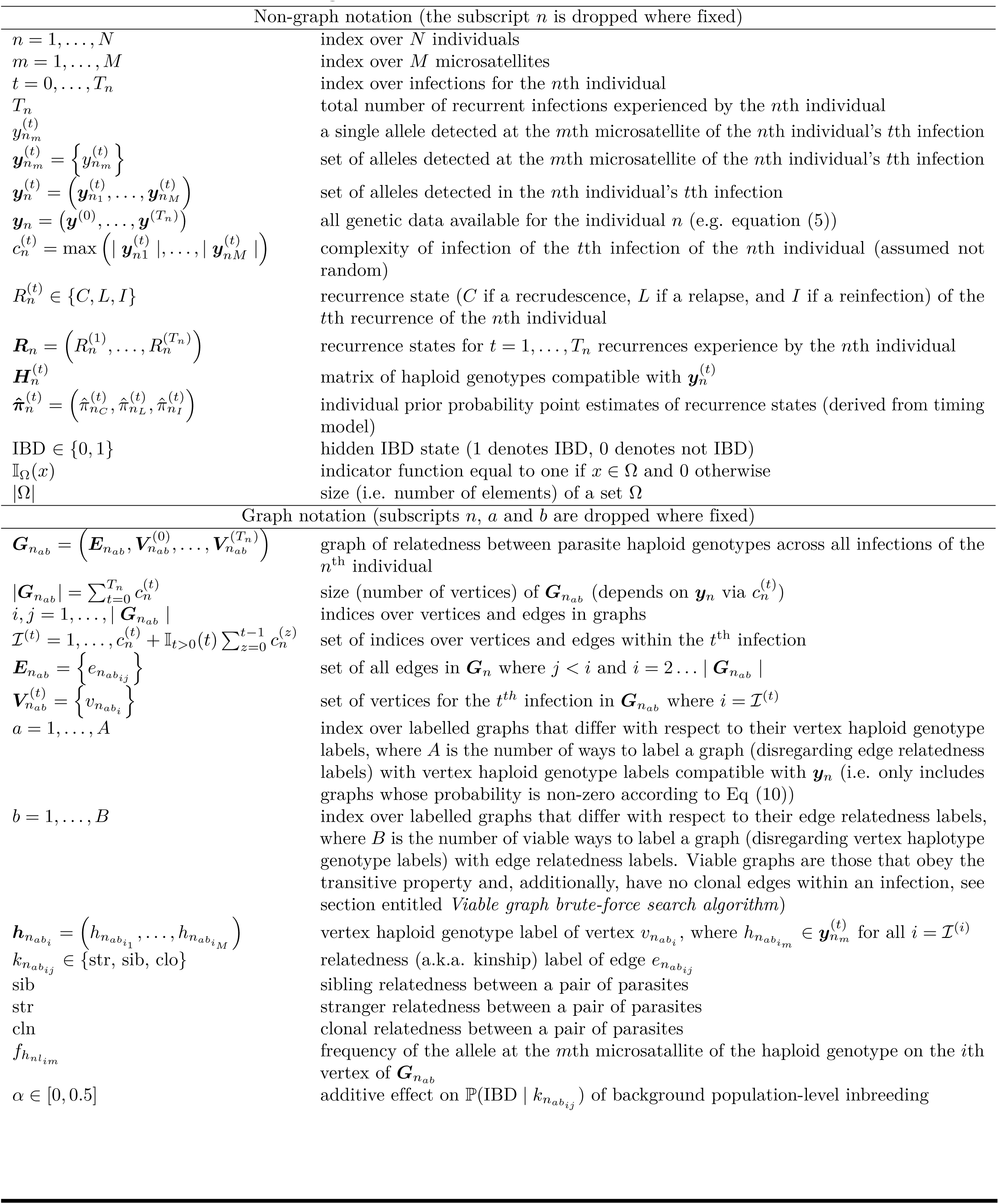
Mathematical notation in genetic model.

### Model overview

This Bayesian model estimates the probability distribution over the possible recurrence states 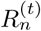, for recurrence *t* experienced by individual n, given all available genetic data for that individual, denoted *y*_*n*_. Note that this includes the genetic data of past episodes and future recurrences. The Bayesian posterior probability of the relapse state *L* for the *t*^*th*^ recurrence experienced by the *n*^*th*^ individual can be written as:

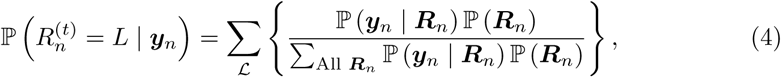

where

- ℙ (***y***_*n*_ | ***R***_*n*_) denotes the likelihood and ℙ (***R***_n_) denotes the prior;
- 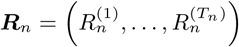 where 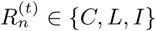 for *t* > 0 *C, L, I* denote recrudescence, relapse and reinfection, respectively, and *T*_*n*_ is the total number of recurrences experienced by the *n*^th^ individual (s.t. ***R***_*n*_ ∈ {*C,L,I*} if *T*_*n*_ = 1 and *Rn* ∈ {*II, LL, CC, IC, CI, IL, LI, LC, CL*} if *T*_*n*_ =2);
- 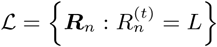 (i.e. ***ℒ*** is the set of all ***R***_*n*_ such that 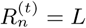);
- 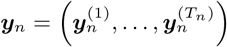 where 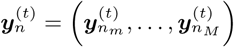 for *t* = *0*,…,*Tn*, and for *m* = 1,…,*M* microsatellites typed, 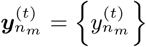 is the set of alleles observed at the *m*^*th*^ microsatellite typed in the t^th^ infection experienced by the n^th^ individual (e.g. 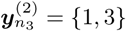 in equation (5)).

**Example per-person set of data, *y*_*n*_**

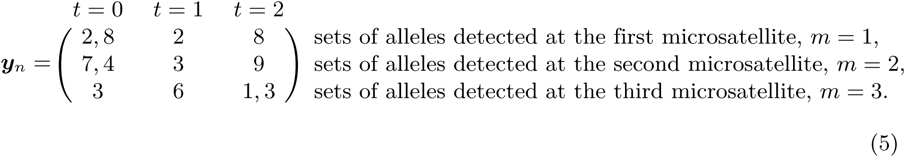

### Evaluation of the prior

The prior, ℙ (***R***_*n*_) where 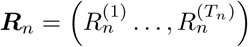 is evaluated by modelling each 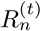 as a random variable from a categorical distribution over *C, L* and *I* with per person recurrence probability point estimates generated under the time-to-event model, namely 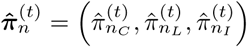, for *t* =*1*,…,*T*_*n*_,

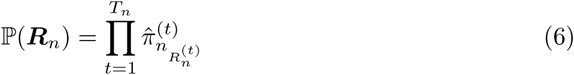

### Evaluation of the likelihood

The likelihood, ℙ (***y***_*n*_ | ***R***_n_), is evaluated by summing over person-specific (indexed by *n*) vertex labelled (indexed by *a*) and edge labelled (indexed by b) graphs of relatedness over parasite haploid genotypes within and across infections, ***G***_*nab*_,

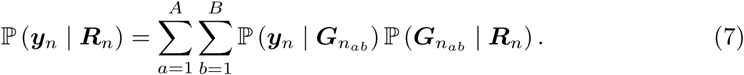

The solution to equation (7) is described in three parts: first we describe the set of fully labelled graphs 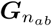, second we describe 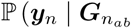, and third we describe 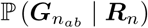. The individual, n, is considered fixed throughout the next sections so the subscript *n* is dropped from 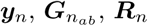 and *T*_*n*_, etc.

### Vertex and edge labelled graphs of parasite relatedness

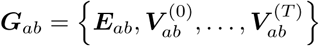 denotes an undirected, edge and vertex labelled, viable graph of relatedness over parasite haploid genotypes within and across *t* = *0*,…,*T* infections for a given individual. *a* is an index over all the possible combinations for labelling the vertices; *b* is an index over all the possible combinations for labelling the edges; viable graphs include only those that obey the transitive property and have no clonal edges within an infection (see section on *Viable graph brute-force search algorithm*). ∀a ∈ *1*‥*A,b* ∈ *1*‥*B*, we define the following:

- 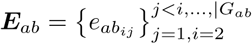, the set of all edges in ***G****ab;*
- 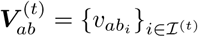, the set of all vertices in *G*_*ab*_ corresponding to the *t*^th^ infection, where 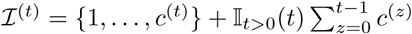 is the set of indices corresponding to the *t*^th^ infection and *c*^(*t*)^ is the COI of the *t*^th^ infection;
- The relatedness (a.k.a. “kinship”) label 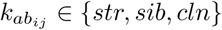, for each edge 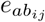, where *str, sib* and *cln* denote stranger, sibling and clone, respectively;
- The haploid genotype label 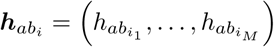 for each vertex, 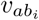, where 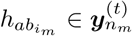, *t* = *0*,…,*T*, and *m* =*1*,…*M*.

Since we assume that the COI of the *t*^th^ infection, *c*^(*t*)^, is equal to the maximum number of distinct alleles seen at any microsatellite within that infection, i.e. 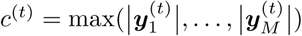, and since we assume no genotyping error nor mutation (assumptions 3 and 4, main text), for ℙ (***y*** | ***G***_*ab*_) > 0 we require that 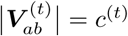 for all *t* (i.e. the number of vertices must equal the COI) and all alleles in *y* need to be represented at least once by the vertex haploid genotype labels. Figure 12 shows two example graphs for the example set of data in equation (5). The graphs in Figure 12 differ in their vertex haploid genotype labels 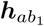 and 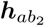, but both have the same set of edge relatedness labels, 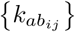 for all *j < i* and *i* = 2,…, | ***G***_*ab*_ |. As mentioned above, we use *a* and *b* to index over vertex haploid genotype labels and edge relatedness labels, respectively, and so the two graphs ***G***_*ab*_ of Figure 12 have the different *a* ∈ {1, 2} but the same *b* =1.

Example graphs compatible with *y* of equation (5).

**Fig 12.**
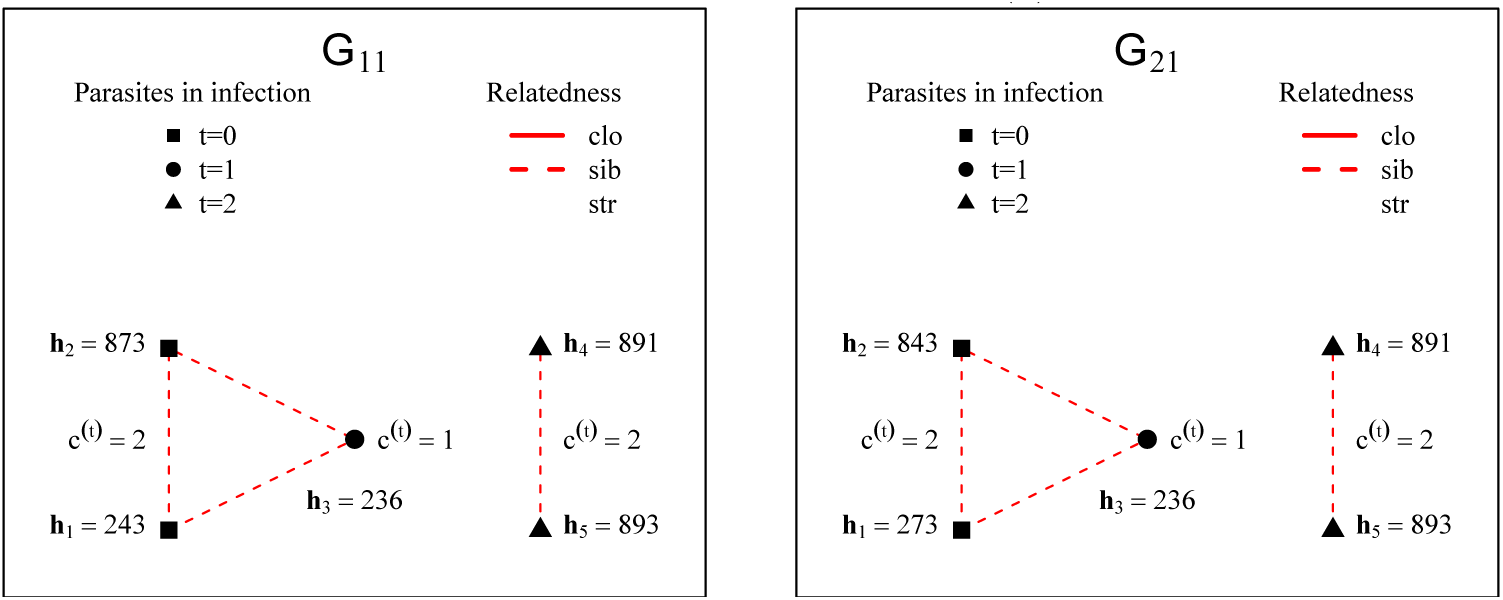
Within each example graph, ***G***_*ab*_ where *a* ∈ {1, 2} and *b* =1, indices *a* and *b* are fixed so dropped hereafter and within each subplot above. The vertices of both graphs are labelled from left to right: ***V***^(0)^ = {*v*_1_,*v*_*2*_} with haploid genotype labels {***h***_1_, ***h***_2_}, ***V***^(1)^ = {*v*_3_} with haploid genotype label ***h***_3_, and ***V***^(2)^ = {*v*_4_,*v*_5_} with haploid genotype labels {***h***_4_, ***h***_5_}. The edge relatedness (a.k.a. kinship) labels of both graphs are the same: *k*_12_ = *sib, k*_23_ = *sib, k*_13_ = *sib* and *k*_45_ = *sib*, while the rest are all *str*, such that *b* = 1 for both graphs. On the contrary, the vertex haploid genotype labels ***h***_1_ and ***h***_2_ differ across the example graphs and so the example graphs have different indices, *a* = 1 and *a* = 2.

The number of ways to allocate haploid genotype labels to a graph for the *n*^th^ individual, *A*, is enumerated independently of edge relatedness labels. *A* depends on both 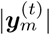 (i.e. the number of alleles detected at a given microsatellite in a given infection) and the number of 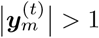 (i.e. the number of heteroallelic microsatellite calls). To see why this is the case, first let ***H***^(*t*)^ be a matrix whose column vectors are haploid genotypes compatible with ***y***^(*t*)^ (e.g. equation (8)).

**Example haploid genotypes compatible with *y*^(0)^ of equation (5)**

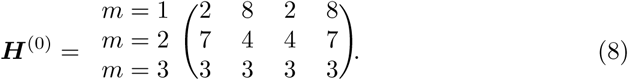

For a given ***y***^(*t*)^, the number of possible haploid genotypes (i.e. number of columns of 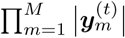. The number of ways to label vertices in ***V***^(*t*)^ is given by the number of ways to choose *c*^(*t*)^ haploid genotypes from the 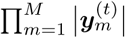 possible haploid genotypes,

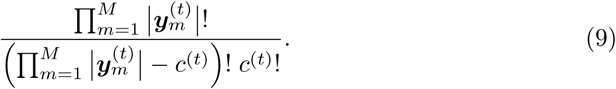

However, of the many combinations given by equation (9),

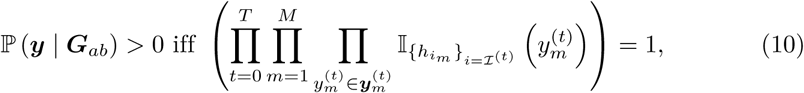

That is, only combinations where all alleles in ***y*** are represented at least once (e.g. those in Figure 12) lead to ℙ (***y*** | ***G***_*ab*_) *>* 0, and contribute to the total number of ways, *A*, to label a graph with vertex haploid genotype labels compatible with *y*. Since *A* is enumerated independently of edge relatedness labels, graphs that have ℙ (***y*** | ***G***_*ab*_) = 0 due to 𝕀_*cln*_ (*k*_*ij*_) = 1 and 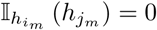 (see below) or NA values do contribute to *A*.

### Probability of the data given a graph

The probability of the data given a vertex and edge labelled graph, ℙ(***y*** | ***G***_*ab*_), is calculated assuming conditional independence between microsatellites and between edges,

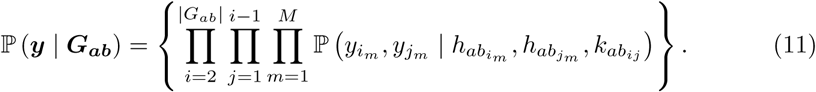

where 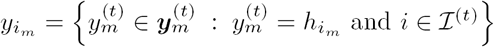. Note that 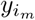 and 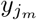 may be within or across infections (i.e. *i* and *j* may be from within the same or across different *ℐ* ^(0)^,…, *ℐ* ^(*T*)^).

Hereafter we consider a single graph with fixed vertex and edge labels thus drop the indices *a* and *b*. To evaluate 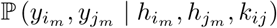 we assume conditional independence between pairs of vertices and edges given the IBD state of the *m*^th^ marker,

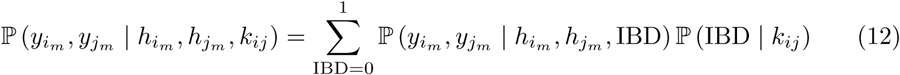

where

- ℙ (IBD = 1 | *k*_*ij*_ = *cln*) = 1; ℙ (IBD = 1 | *sib*) = 0.5 + *α;* ℙ (IBD = 1 | *str*) = 0 + *α*;
- ℙ (IBD = 0 | *k*_*ij*_) = 1 - ℙ (IBD = 1 | *k*_*ij*_);
- *α* is an additive effect of background population-level inbreeding;
- 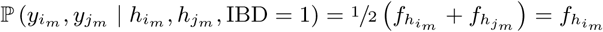 if 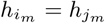 and 0 otherwise;
- 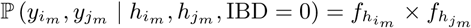
- 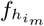 and 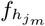 denote the frequency of the allele at the m^th^ microsatellite of the haploid genotype label of the i^th^ and j^th^ vertex, respectively. Equivalently, we could write 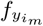 and 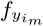, respectively.

Together, the above lead to

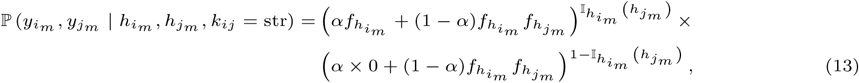

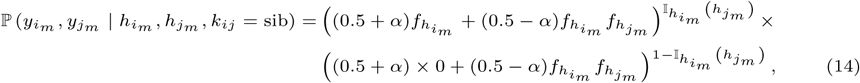

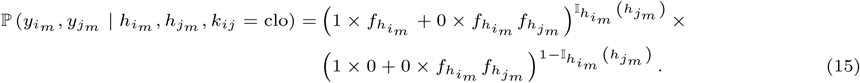

### Probability of a graph given a series of recurrence states

To calculate the probability of a graph given a series of recurrence states, we assume independence between recurrence states,

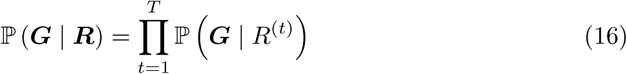

Where

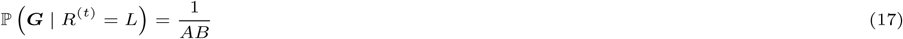

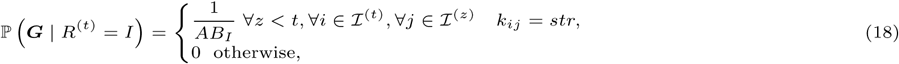

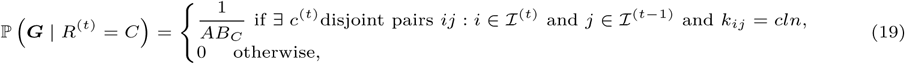

where *B*_*I*_ *< B* and *B*_*C*_ *< B* denote the number of graphs that satisfy the conditions outlined in Eq (18) and (19), both of which are determined algorithmically from the adjacency matrix of G.

The condition outlined in Eq (19) that *j*∈ℐ^(*t*–1)^ specifies that a recrudescence is seeded by the most recent past infection only (assumption 2). Also in Eq (19), the condition that there are c^(*t*)^ disjoint pairs follows from assumptions 5 and 6 and results in zero probability of recrudescence following an infection with lower COI (i.e. a recrudescence cannot be more diverse that the infection that seeded it - diversity cannot be be created, only lost). For example, in Eq (5), 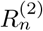 has zero probability of being a recrudescence because *c*^(2)^ =2 > *c*^(1)^ = 1.

Presently, Eq (17) to (19) do not take into the relative likelihood of parasites that are strangers, siblings or clones in relation to one another within an inoculation. However, they could be adapted to do so (e.g. by coupling to a transmission mode). Note that ℙ(G | R) implicitly conditions on *y* via *c*^(*t*)^.

### Model implementation

Above, *t* = *0*,…,*T*_*n*_ where *T*_*n*_ is the number of recurrences experienced by the *n*^th^ individual. In the code, *t* = *1*,…,*T*_*n*_ where *T*_*n*_ is the number of infections experienced by the *n*^th^ individual.

The model is implemented on the log scale to prevent under and over flow problems, using the log sum exp trick where appropriate. Instead of summing over *a* and *b* in one step as Eq (7) implies, we first sum over graphs indexed by *a* = 1,…,*A* fixing *b* and working entirely on the log domain (interior of square brackets Eq (20)); we then sum over graphs indexed by 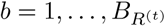 where 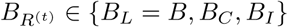 is determined algorithmically from the adjacency matrix of ***G***,

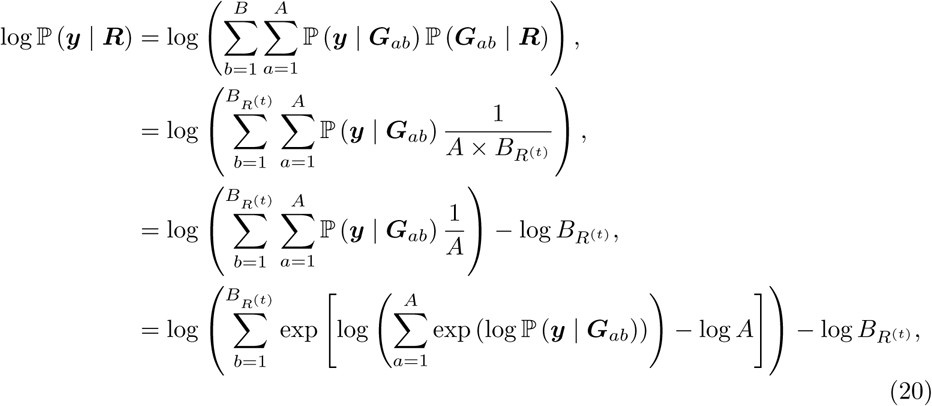

where the subscript *n* is dropped since *n* is fixed.

### Viable graph brute-force search algorithm

For a given set of complexities 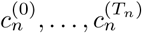 we implement a brute-force algorithm that searches over all graphs and stores all viable graphs. Viable graphs can be described with two independent rules:

a. No clonal edges between vertices within an infection
b. All connected paths must obey the transitivity property

The algorithm is summarized as follows:

1. Construct a list *S*_*G*_ of all graphs by listing all adjacency matrices described by block matrices (one per infection) for which the block matrices only contain {0, 0.5}, and the across blocks contain {0, 0.5,1} (this implies no clonal edges within infections).
2. For each *G* ∈ 𝒮_*G*_:
  - Enumerate all connected components in *G* and verify that each connected component is a clique (fully connected subgraph).
  - List all triangular cliques (fully connected subgraphs containing exactly three vertices)
  - For each triangular clique *G′* compute the sum of the weighted edges: If the sum of the edges in *G′* is equal to 2.5 then **Reject** *G* else **Accept** the subgraph *G′*
3. If all subgraphs are accepted, **Accept** *G*

Step 2 results in obeying rule (b) whereby graphs with non-transitive relatedness patterns (e.g. A is clonal with both B & C but B is sibling with C) are rejected.

## References

1. Howes RE, Battle KE, Mendis KN, Smith DL, Cibulskis RE, Baird JK, et al. Global Epidemiology of Plasmodium vivax. The American journal of tropical medicine and hygiene. 2016;95(6 Suppl):15–34. doi:10.4269/ajtmh.16-0141.

2. World Health Organization. World Malaria Report 2017. Geneva; 2017.

3. Robinson LJ, Wampfler R, Betuela I, Karl S, White MT, Suen CSLW, et al. Strategies for understanding and reducing the Plasmodium vivax and Plasmodium ovale hypnozoite reservoir in Papua New Guinean children: a randomised placebo-controlled trial and mathematical model. PLoS medicine. 2015;12(10):e1001891.

4. Chu CS, Phyo AP, Lwin KM, Win HH, San T, Aung AA, et al. Comparison of the Cumulative Efficacy and Safety of Chloroquine, Artesunate, and Chloroquine-Primaquine in Plasmodium vivax Malaria. Clinical Infectious Diseases. 2018; p. ciy319. doi:10.1093/cid/ciy319.

5. Coatney GR, Cooper WC, Young MD. Studies in Human Malaria. XXX. A Summary of 204 Sporozoite-Indueed Infections with the Chesson Strain of Plasmodium vivax. Journal of the National Malaria Society. 1950;9(4):381–96.

6. White NJ. Determinants of relapse periodicity in Plasmodium vivax malaria. Malaria Journal. 2011;10(1):297. doi:10.1186/1475-2875-10-297.

7. Battle KE, Karhunen MS, Bhatt S, Gething PW, Howes RE, Golding N, et al. Geographical variation in Plasmodium vivax relapse. Malar J. 2014;13:144.

8. Gonzalez-Ceron L, Mu J, Santillan F, Joy D, Sandoval MA, Camas G, et al. Molecular and epidemiological characterization of Plasmodium vivax recurrent infections in southern Mexico. Parasit Vectors. 2013;6:109.

9. Kim JR, Nandy A, Maji AK, Addy M, Dondorp AM, Day NPJ, et al. Genotyping of Plasmodium vivax reveals both short and long latency relapse patterns in Kolkata. PloS one. 2012;7(7):e39645. doi:10.1371/journal.pone.0039645.

10. Joshi H, Prajapati SK, Verma A, Kang’a S, Carlton JM. Plasmodium vivax in India. Trends Parasitol. 2008;24(5):228–235.

11. Imwong M, Boel ME, Pagornrat W, Pimanpanarak M, McGready R, Day NP, et al. The first Plasmodium vivax relapses of life are usually genetically homologous. J Infect Dis. 2012;205(4):680–683.

12. Imwong M, Snounou G, Pukrittayakamee S, Tanomsing N, Kim JR, Nandy A, et al. Relapses of Plasmodium vivax Infection Usually Result from Activation of Heterologous Hypnozoites. The Journal of Infectious Diseases. 2007;195:927–33. doi:10.1086/512241.

13. Chen N, Auliff A, Rieckmann K, Cheng Q. Relapses of Plasmodium vivax infection result from clonal hypnozoites activated at predetermined intervals. The Journal of infectious diseases. 2007;195(7):934–941.

14. Orjuela-Sanchez P, da Silva NS, da Silva-Nunes M, Ferreira MU. Recurrent parasitemias and population dynamics of Plasmodium vivax polymorphisms in rural Amazonia. Am J Trop Med Hyg. 2009;81(6):961–968.

15. Restrepo E, Imwong M, Rojas W, Carmona-Fonseca J, Maestre A. High genetic polymorphism of relapsing P. vivax isolates in northwest Colombia. Acta Trop. 2011;119(1):23–29.

16. de Araujo FC, de Rezende AM, Fontes CJ, Carvalho LH, Alves de Brito CF. Multiple-clone activation of hypnozoites is the leading cause of relapse in Plasmodium vivax infection. PLoS ONE. 2012;7(11):e49871.

17. Maneerattanasak S, Gosi P, Krudsood S, Chimma P, Tongshoob J, Mahakunkijcharoen Y, et al. Molecular and immunological analyses of confirmed Plasmodium vivax relapse episodes. Malar J. 2017;16(1):228.

18. Popovici J, Friedrich LR, Kim S, Bin S, Run V, Lek D, et al. Genomic analyses reveal the common occurrence and complexity of Plasmodium vivax relapses in Cambodia. mBio. 2018;9(1):e01888–17.

19. Recht J, Ashley E, White N, Organization WH, et al. Safety of 8-aminoquinoline antimalarial medicines. World Health Organization; 2014.

20. Betuela I, Rosanas-Urgell A, Kiniboro B, Stanisic DI, Samol L, de Lazzari E, et al. Relapses Contribute Significantly to the Risk of Plasmodium vivax Infection and Disease in Papua New Guinean Children 1–5 Years of Age. The Journal of Infectious Diseases. 2012;206(11):1771–1780. doi:10.1093/infdis/jis580.

21. Baton LA, Ranford-Cartwright LC. Spreading the seeds of million-murdering death: Metamorphoses of malaria in the mosquito. Trends in Parasitology. 2005;21(12):573–580. doi:10.1016/j.pt.2005.09.012.

22. Baker DA. Malaria gametocytogenesis. Molecular and Biochemical Parasitology. 2010;172(2):57–65. doi:10.1016/j.molbiopara.2010.03.019.

23. Abreha T, Hwang J, Thriemer K, Tadesse Y, Girma S, Melaku Z, et al. Comparison of artemether-lumefantrine and chloroquine with and without primaquine for the treatment of Plasmodium vivax infection in Ethiopia: A randomized controlled trial. PLoS Medicine. 2017;14(5):1–17.

24. Veron V, Legrand E, Yrinesi J, Volney B, Simon S, Carme B. Genetic diversity of msp3alpha and msp1b5 markers of Plasmodium vivax in French Guiana. Malar J. 2009;8:40.

25. McCollum AM, Soberon V, Salas CJ, Santolalla ML, Udhayakumar V, Escalante AA, et al. Genetic variation and recurrent parasitaemia in Peruvian Plasmodium vivax populations. Malar J. 2014;13:67.

26. Cowell AN, Valdivia HO, Bishop DK, Winzeler EA. Exploration of Plasmodium vivax transmission dynamics and recurrent infections in the Peruvian Amazon using whole genome sequencing. Genome Medicine. 2018;10(1):52. doi:10.1186/s13073-018-0563-0.

27. White MT, Shirreff G, Karl S, Ghani AC, Mueller I. Variation in relapse frequency and the transmission potential of Plasmodium vivax malaria. Proc R Soc B. 2016;283(1827):20160048.

28. Popovici J, Pierce-Friedrich L, Kim S, Bin S, Run V, Lek D, et al. Recrudescence, reinfection or relapse? A more rigorous framework to assess chloroquine efficacy for vivax malaria. The Journal of Infectious Diseases. 2018; p. jiy484. doi:10.1093/infdis/jiy484.

29. WHO, Medicines for Malaria Venture. Methods and techniques for clinical trials on antimalarial drug efficacy: genotyping to identify parasite populations: informal consultation organized by the Medicines for Malaria Venture and cosponsored by the World Health Organization.; 2008.

30. Messerli C, Hofmann NE, Beck HP, Felger I. Critical evaluation of molecular monitoring in malaria drug efficacy trials: pitfalls of length polymorphic markers. Antimicrobial Agents and Chemotherapy. 2016;61(1):AAC.01500–16. doi:10.1128/AAC.01500-16.

31. Mwingira F, Nkwengulila G, Schoepflin S, Sumari D, Beck HP, Snounou G, et al. Plasmodium falciparum msp1, msp2 and glurp allele frequency and diversity in sub-Saharan Africa. Malaria Journal. 2011;10:1–10. doi:10.1186/1475-2875-10-79.

32. Chu CS, Phyo AP, Turner C, Win HH, Poe NP, Yotyingaphiram W, et al. Chloroquine Versus Dihydroartemisinin-Piperaquine With Standard High-dose Primaquine Given Either for 7 Days or 14 Days in Plasmodium vivax Malaria. Clinical Infectious Diseases. 2018; p. ciy735. doi:10.1093/cid/ciy735.

33. Adekunle AI, Pinkevych M, McGready R, Luxemburger C, White LJ, Nosten F, et al. Modeling the dynamics of Plasmodium vivax infection and hypnozoite reactivation in vivo. PLoS neglected tropical diseases. 2015;9(3):e0003595.

34. White MT, Karl S, Battle KE, Hay SI, Mueller I, Ghani AC. Modelling the contribution of the hypnozoite reservoir to Plasmodium vivax transmission. Elife. 2014;3.

35. White MT, Walker P, Karl S, Hetzel MW, Freeman T, Waltmann A, et al. Mathematical modelling of the impact of expanding levels of malaria control interventions on Plasmodium vivax. Nature communications. 2018;9(1):3300.

36. Landier J, Parker DM, Thu AM, Lwin KM, Delmas G, Nosten FH, et al. Effect of generalised access to early diagnosis and treatment and targeted mass drug administration on Plasmodium falciparum malaria in Eastern Myanmar: an observational study of a regional elimination programme. Lancet (London, England). 2018;391(10133):1916–1926. doi:10.1016/S0140-6736(18)30792-X.

37. White MT, Karl S, Koepfli C, Longley RJ, Hofmann NE, Wampfler R, et al. Plasmodium vivax and Plasmodium falciparum infection dynamics: re-infections, recrudescences and relapses. Malaria journal. 2018;17(1):170.

38. Watson J, Chu CS, Tarning J, White NJ. Characterizing Blood-Stage Antimalarial Drug MIC Values In Vivo Using Reinfection Patterns. Antimicrob Agents Chemother. 2018;62(7).

39. Golassa L, White MT. Population-level estimates of the proportion of Plasmodium vivax blood-stage infections attributable to relapses among febrile patients attending Adama Malaria Diagnostic Centre, East Shoa Zone, Oromia, Ethiopia. Malar J. 2017;16(1):301.

40. Santos-Vega M, Bouma MJ, Kohli V, Pascual M. Population Density, Climate Variables and Poverty Synergistically Structure Spatial Risk in Urban Malaria in India. PLoS Negl Trop Dis. 2016;10(12):e0005155.

41. Lover AA, Coker RJ. Do mixed infections matter? Assessing virulence of mixed-clone infections in experimental human and murine malaria. Infect Genet Evol. 2015;36:82–91.

42. Tenero D, Green JA, Goyal N. Exposure-Response Analyses for Tafenoquine after Administration to Patients with Plasmodium vivax Malaria. Antimicrob Agents Chemother. 2015;59(10):6188–6194.

43. Karl S, Laman M, Moore BR, Benjamin J, Koleala T, Ibam C, et al. Gametocyte Clearance Kinetics Determined by Quantitative Magnetic Fractionation in Melanesian Children with Uncomplicated Malaria Treated with Artemisinin Combination Therapy. Antimicrob Agents Chemother. 2015;59(8):4489–4496.

44. Battle KE, Cameron E, Guerra CA, Golding N, Duda KA, Howes RE, et al. Defining the relationship between Plasmodium vivax parasite rate and clinical disease. Malar J. 2015;14:191.

45. Kerlin DH, Gatton ML. A simulation model of the within-host dynamics of Plasmodium vivax infection. Malar J. 2015;14:51.

46. Lover AA, Zhao X, Gao Z, Coker RJ, Cook AR. The distribution of incubation and relapse times in experimental human infections with the malaria parasite Plasmodium vivax. BMC Infect Dis. 2014;14:539.

47. Roy M, Bouma MJ, Ionides EL, Dhiman RC, Pascual M. The potential elimination of Plasmodium vivax malaria by relapse treatment: insights from a transmission model and surveillance data from NW India. PLoS Negl Trop Dis. 2013;7(1):e1979.

48. Chamchod F, Beier JC. Modeling Plasmodium vivax: relapses, treatment, seasonality, and G6PD deficiency. J Theor Biol. 2013;316:25–34.

49. Gething PW, Elyazar IR, Moyes CL, Smith DL, Battle KE, Guerra CA, et al. A long neglected world malaria map: Plasmodium vivax endemicity in 2010. PLoS Negl Trop Dis. 2012;6(9):e1814.

50. Elyazar IR, Gething PW, Patil AP, Rogayah H, Sariwati E, Palupi NW, et al. Plasmodium vivax malaria endemicity in Indonesia in 2010. PLoS ONE. 2012;7(5):e37325.

51. Aguas R, Ferreira MU, Gomes MG. Modeling the effects of relapse in the transmission dynamics of malaria parasites. J Parasitol Res. 2012;2012:921715.

52. Ishikawa H, Ishii A, Nagai N, Ohmae H, Harada M, Suguri S, et al. A mathematical model for the transmission of Plasmodium vivax malaria. Parasitol Int. 2003;52(1):81–93.

53. Kammanee A, Kanyamee N, Tang IM. Basic reproduction number for the transmission of Plasmodium vivax malaria. Southeast Asian J Trop Med Public Health. 2001;32(4):702–706.

54. Mason DP. Review and recent progress: the mathematical modeling of mixed species Plasmodium infections. Southeast Asian J Trop Med Public Health. 2000;31 Suppl 1:69–74.

55. Ross A, Koepfli C, Schoepflin S, Timinao L, Siba P, Smith T, et al. The incidence and differential seasonal patterns of Plasmodium vivax primary infections and relapses in a cohort of children in Papua New Guinea. PLoS neglected tropical diseases. 2016;10(5):e0004582.

56. Zhu SJ, Hendry JA, Almagro-Garcia J, Pearson RD, Amato R, Miles A, et al. The origins and relatedness structure of mixed infections vary with local prevalence of P. falciparum malaria. bioRxiv. 2018;doi:10.1101/387266.

57. Barry AE, Waltmann A, Koepfli C, Barnadas C, Mueller I. Uncovering the transmission dynamics of Plasmodium vivax using population genetics. Pathog Glob Health. 2015;109(3):142–152.

58. Pearson RD, Amato R, Auburn S, Miotto O, Almagro-Garcia J, Amaratunga C, et al. Genomic analysis of local variation and recent evolution in Plasmodium vivax. Nature genetics. 2016;48(8):959–964. doi:10.1038/ng.3599.

59. Hupalo DN, Luo Z, Melnikov A, Sutton PL, Rogov P, Escalante A, et al. Population genomics studies identify signatures of global dispersal and drug resistance in Plasmodium vivax. Nature Genetics. 2016;48(8):953–958. doi:10.1038/ng.3588.

60. Wang J. Sibship Reconstruction from Genetic Data with Typing Errors. Genetics. 2004;166(4):1963–1979. doi:10.1534/genetics.166.4.1963.

61. Wang J, Scribner KT. Parentage and sibship inference from markers in polyploids. Molecular Ecology Resources. 2014;14(3):541–553. doi:10.1111/1755-0998.12210.

62. Rosenberg NA, Li LM, Ward R, Pritchard JK. Informativeness of Genetic Markers for Inference of Ancestry *. Am J Hum Genet. 2003;73:1402–1422.

63. Weir BS, Anderson AD, Hepler AB. Genetic relatedness analysis: Modern data and new challenges. Nature Reviews Genetics. 2006;7(10):771–780. doi:10.1038/nrg1960.

64. Anderson EC, Garza JC. The power of single-nucleotide polymorphisms for large-scale parentage inference. Genetics. 2006;172(4):2567–2582. doi:10.1534/genetics.105.048074.

65. Bright AT, Manary MJ, Tewhey R, Arango EM, Wang T, Schork NJ, et al. A high resolution case study of a patient with recurrent Plasmodium vivax infections shows that relapses were caused by meiotic siblings. PLoS neglected tropical diseases. 2014;8(6):e2882.

66. Gattepaille LM, Jakobsson M. Combining markers into haplotypes can improve population structure inference. Genetics. 2012;190(1):159–174. doi:10.1534/genetics.111.131136.

67. Baetscher DS, Clemento AJ, Ng TC, Anderson EC, Garza JC. Microhaplotypes provide increased power from short-read DNA sequences for relationship inference. Molecular Ecology Resources. 2018;18(2):296–305. doi:10.1111/1755-0998.12737.

68. Carrara VI, Hogan C, De Pree C, Nosten F, McGready R. Improved pregnancy outcome in refugees and migrants despite low literacy on the Thai-Burmese border: results of three cross-sectional surveys. BMC Pregnancy and Childbirth. 2011;11(1):45. doi:10.1186/1471-2393-11-45.

69. Gunawardena S, Karunaweera ND, Ferreira MU, Phone-Kyaw M, Pollack RJ, Alifrangis M, et al. Geographic structure of Plasmodium vivax: microsatellite analysis of parasite populations from Sri Lanka, Myanmar, and Ethiopia. The American journal of tropical medicine and hygiene. 2010;82(2):235–242.

70. Csardi G, Nepusz T. The igraph software package for complex network research. InterJournal. 2006;Complex Systems:1695.

71. Stan Development Team. RStan: the R interface to Stan; 2018. Available from: http://mc-stan.org/.

72. Carpenter B, Gelman A, Hoffman MD, Lee D, Goodrich B, Betancourt M, et al. Stan: A probabilistic programming language. Journal of statistical software. 2017;76(1).

73. Taylor AR, Flegg JA, Nsobya SL, Yeka A, Kamya MR, Rosenthal PJ, et al. Estimation of malaria haplotype and genotype frequencies: A statistical approach to overcome the challenge associated with multiclonal infections. Malaria Journal. 2014;13(1):1–11. doi:10.1186/1475-2875-13-102.

74. Matthews H, Duffy CW, Merrick CJ. Checks and balances? DNA replication and the cell cycle in Plasmodium. Parasites & vectors. 2018;11(1):216. doi:10.1186/s13071-018-2800-1.

75. Bink MC, Anderson AD, Van De Weg WE, Thompson EA. Comparison of marker-based pairwise relatedness estimators on a pedigreed plant population. Theoretical and Applied Genetics. 2008;117(6):843–855. doi:10.1007/s00122-008-0824-1.

76. Zhu SJ, Hendry JA, Almagro-garcia J, Pearson RD, Amato R, Miles A, et al. The origins and relatedness structure of mixed infections vary with local prevalence of P. falciparum malaria. bioRxiv. 2018;.

77. Soontarawirat I, Andolina C, Paul R, Day NPJ, Nosten F, Woodrow CJ, et al. Plasmodium vivax genetic diversity and heterozygosity in blood samples and resulting oocysts at the Thai-Myanmar border. Malar J. 2017;16(1):355.

78. Schaffner SF, Taylor AR, Wong W, Wirth DF, Neafsey DE. HmmIBD: Software to infer pairwise identity by descent between haploid genotypes. Malaria Journal. 2018;17(1):10–13. doi:10.1186/s12936-018-2349-7.

79. Henden L, Lee S, Mueller I, Barry A, Bahlo M. Identity-by-descent analyses for measuring population dynamics and selection in recombining pathogens. vol. 14; 2018.

80. Lin JT, Patel JC, Kharabora O, Sattabongkot J, Muth S, Ubalee R, et al. Plasmodium vivax isolates from Cambodia and Thailand show high genetic complexity and distinct patterns of P. vivax multidrug resistance gene 1 (pvmdr1) polymorphisms. Am J Trop Med Hyg. 2013;88(6):1116–1123.

81. Jacob PE, Murray LM, Holmes CC, Robert CP. Better together? Statistical learning in models made of modules. arXiv preprint arXiv:170808719. 2017;.

82. Wong W, Wenger EA, Hartl DL, Wirth DF. Modeling the genetic relatedness of Plasmodium falciparum parasites following meiotic recombination and cotransmission. PLOS Computational Biology. 2018;14(1):e1005923. doi:10.1371/journal.pcbi.1005923.

83. Lin JT, Hathaway NJ, Saunders DL, Lon C, Balasubramanian S, Kharabora O, et al. Using amplicon deep sequencing to detect genetic signatures of Plasmodium vivax relapse. The Journal of infectious diseases. 2015;212(6):999–1008.

